# Phosphorylation dynamics in a flg22-induced, heterotrimeric G-protein dependent signaling network in *Arabidopsis thaliana* reveals a candidate PP2A phosphatase involved in AtRGS1 trafficking

**DOI:** 10.1101/2021.12.06.471472

**Authors:** Justin M. Watkins, Natalie M. Clark, Gaoyuan Song, Celio Cabral Oliveira, Bharat Mishra, Libuse Brachova, Clara M. Seifert, Malek S. Mitchell, Pedro Augusto Braga dos Reis, Daisuke Urano, M. Shahid Muktar, Justin W. Walley, Alan M. Jones

## Abstract

flg22 is a 22-amino peptide released from bacterial flagellin, a Microbe-Associated Molecular Pattern (MAMP) that is recognized by the plant cell as a signal indicating that bacteria are present. On its own, flg22 initiates a rapid increase in cytoplasmic calcium, extracellular reactive oxygen species, and activation of a Mitogen Activated Protein Kinase (MAPK) cascade, all of which are activated within 15 minutes after the cell perceives flg22. Here we show a massive change in protein abundance and phosphorylation state of the Arabidopsis root cell proteome within this 15-minute duration in wildtype and a mutant deficient in G-protein coupled signaling. Integration of phosphoproteome with protein-protein interactome data followed by network topology analyses discovered that many of the flg22-induced phosphoproteome changes fall on proteins that comprise the G protein interactome and on the most highly populated hubs of the immunity network. Approximately 95% of the phosphorylation changes in the G-protein interactome depend on a functional heterotrimeric G protein complex, some occur on proteins that interact directly with components of G-coupled signal transduction. One of these is ATBα, a substrate-recognition subunit of the PP2A Ser/Thr phosphatase and an interactor to *Arabidopsis thaliana* REGULATOR OF G SIGNALING 1 protein (AtRGS1), a 7-transmembrane spanning modulator of the nucleotide-binding state of the core G protein complex. AtRGS1 is phosphorylated by BAK1, a component of the flg22 receptor, to initiate AtRGS1 endocytosis. A null mutation of *ATB*α confers high basal endocytosis of AtRGS1, suggesting sustained phosphorylated status. Loss of ATBα confers traits associated with loss of AtRGS1. Because the basal level of AtRGS1 is lower in the *atbα* null mutant in a proteasome-dependent manner, we propose that phosphorylation-dependent endocytosis of AtRGS1 is part of a mechanism to degrade AtRGS1 which then sustains activation of the G protein complex. Thus, the role of ATBα is now established as a central component of phosphorylation-dependent regulation of system dynamics in innate immunity.

## Introduction

A coordinated signaling relay begins with the perception of external signals by membrane bound receptors, such as G-Protein Coupled Receptors (GPCRs) in animal cells. Similar to animals, the G protein complex in Arabidopsis contains one canonical Gα subunit (AtGPA1), one Gβ subunit (AGB1), one of three Gγ subunits (AGG1, AGG2 and AGG3) (Thung *et al*, 2012; Wolfenstetter *et al*, 2015). However, plant G protein releases GDP, binds GTP and undergoes a conformational change to be active without GPCRs; rather, the activation status is modulated by REGULATOR OF g SIGNALING 1 (AtRGS1) (Jones *et al*, 2011). AtRGS1 contains a 7TM domain, an RGS domain that catalyzes the intrinsic GTP hydrolysis activity of Gα (Jones *et al*, 2012, 2011; Johnston *et al*, 2007; Chen *et al*, 2003), and a C-terminal cluster of phosphorylation sites (Urano *et al*, 2012a; Tunc-Ozdemir & Jones, 2017; Xue *et al*, 2020). AtGPA1 is phosphorylated leading to deactivation (Li *et al*, 2018). In addition, de-repression occurs when AtRGS1 internalizes away from AtGPA1 upon signal perception, and this endocytosis depends on phosphorylation of a phosphocluster in the C-terminal tail. The kinases that phosphorylate AtRGS1 include WITH NO LYSINE (WNK) kinases and BAK1 (Urano *et al*, 2012a; Fu *et al*, 2014a). BAK1 is a component of the flg22 receptor, FLS2 (Sun *et al*, 2013). flg22 is a Microbe-Associated Molecular Patter (MAMP) that induces AtRGS1 endocytosis by phosphorylation within the phosphocluster (Tunc-Ozdemir & Jones, 2017; Tunc-Ozdemir *et al*, 2016; Liang *et al*, 2018) (Watkins *et al*, 2021).

The Arabidopsis G protein interactome project identified over 400 direct interactions within ∼70 highly-interconnected core components (Klopffleisch *et al*, 2011a). Those interactomes combined with further characterizations with biochemical, genetic and cell biological evidence prove new regulators downstream of G protein complex such as the WNK kinases. Through *in vitro* screening of 70 receptor-kinases, we also identified that BAK1 directly phosphorylates AtRGS1 and AtGPA1, as well as their phosphorylation sites and the structural mechanisms of how their activity is regulated (Urano *et al*, 2012b; Watkins *et al*, 2021; Fu *et al*, 2014a; Li *et al*, 2018; Liao *et al*, 2017). Those large-scale screenings revealed direct but static interactions within the plant G protein network that are composed of highly conserved-core nodes (G protein subunits and RGS1) throughout eukaryotes, together with peripheral nodes (other regulators and effectors) largely different from animals. Advanced proteomics approaches to quantitate post-translational modifications, such as phosphorylation, can reveal a dynamic signaling flow over time (Altelaar *et al*, 2013) [ref, general review]. Such time-dependent changes of post-translational status, combined with physical interaction map, allow us to infer direct signaling transmission events between proteins. The two individual phosphorylation targets (AtRGS and AtGPA) are the key molecular signatures controlling G protein activity (Ghusinga *et al*, 2021a).

Pathogen and MAMP-induced quantitative and system-wide experiments yielded many multi-dimensional datasets. To better understand the intricate nature of plant immune signaling cascade in response to such biotic stresses, an integrative network science framework was instrumental to decipher the significant players in plant-pathogen interactions (Mishra *et al*, 2021a, 2019). For example, network architectural or centrality analyses of global protein-protein interaction (PPI) networks (i.e. interactomes) revealed the preferred pathogenic contact points to host (Ahmed *et al*, 2018a; Arabidopsis Interactome Mapping Consortium, 2011a; Mishra *et al*, 2017a; Mukhtar *et al*, 2011a). With reference to phosphoproteomes, their integration with interactomes datasets shows promise for yielding a comprehensive landscape of immune signaling (Mishra *et al*, 2021a, 2019). This integration is particularly relevant for plant-pathogen interactomes that encompass nodes as host or pathogen molecules, while edges exhibit interactions as coordinated by the system-wide host responses. These interactions highlight novel players that induce immune or defense responses under pathogen infection (Mishra *et al*, 2021a).

Using quantitative proteomics, we report flg22-dependent remodeling of the phosphoproteome that highlights G protein-dependent phosphorylation changes within the immunity network. Analysis of this dataset provides key dynamic phosphorylation changes at early time points in the flg22 pathway while also providing a broader picture of overarching changes made in the root in response to flg22. In conjunction with cell biology and biochemical validation, this report reveals a phosphatase of AtRGS1 that plays an integral role in flg22-induced AtRGS1 phosphorylation, endocytosis, and degradation, and, therefore, a key player in G signaling activation and dynamics. While various kinases that phosphorylate AtRGS1 and G protein subunits have been identified, to date, nothing has been known of the phosphatases in this phosphorylation-dephosphorylation cycle. Plant cells possibly utilize kinases to sort out and connect various inputs of extracellular stimuli to G protein complex, while a small number of phosphatases to modulate the activity levels through the relocation of AtRGS1.

## Results

### Dose response and time dynamics of flg22-induced ROS bursts in roots

Our strategy was to use quantitative proteomics in root cells to elucidate changes in the phosphorylation state caused by G protein-dependent flg22 signaling. This first required establishing the kinetics of flg22 signaling in wild type (WT) roots to determine the most informative dose and time for the flg22 response profiled in subsequent experiments. Thus, we employed confocal microscopy to visualize peak flg22 signaling in roots using reactive oxygen species (ROS) bursts as a marker for flg22 signaling. To visualize ROS, *Arabidopsis* roots were treated with the ROS sensor chloromethyl 2′,7′-dihydrodichlorofluorescein diacetate (DCF). The detected intracellular ROS distribution was highest in the elongation zone (**Fig 1A**, red bracket). DCF fluorescence was enhanced in Col-0 roots treated with 50 nM flg22 compared to those treated with water (**Fig 1B**, cf. left panel to middle panel), whereas no increase in fluorescence was observed in flg22-treated *fls2-1*, a null mutant of the flg22 receptor FLS2 (Fig 1B, cf. middle panel to right panel). The flg22-induced ROS burst dose response reached a saturation point near 50 nM (**Fig 1C**). To determine the optimal treatment time for our phosphoproteomics experiment, we treated roots with 50 nM flg22 for 0, 5, 10, 15, and 30 minutes. Peak DCF fluorescence was observed at 15 minutes (**Fig 1D**).

**Figure 1.**
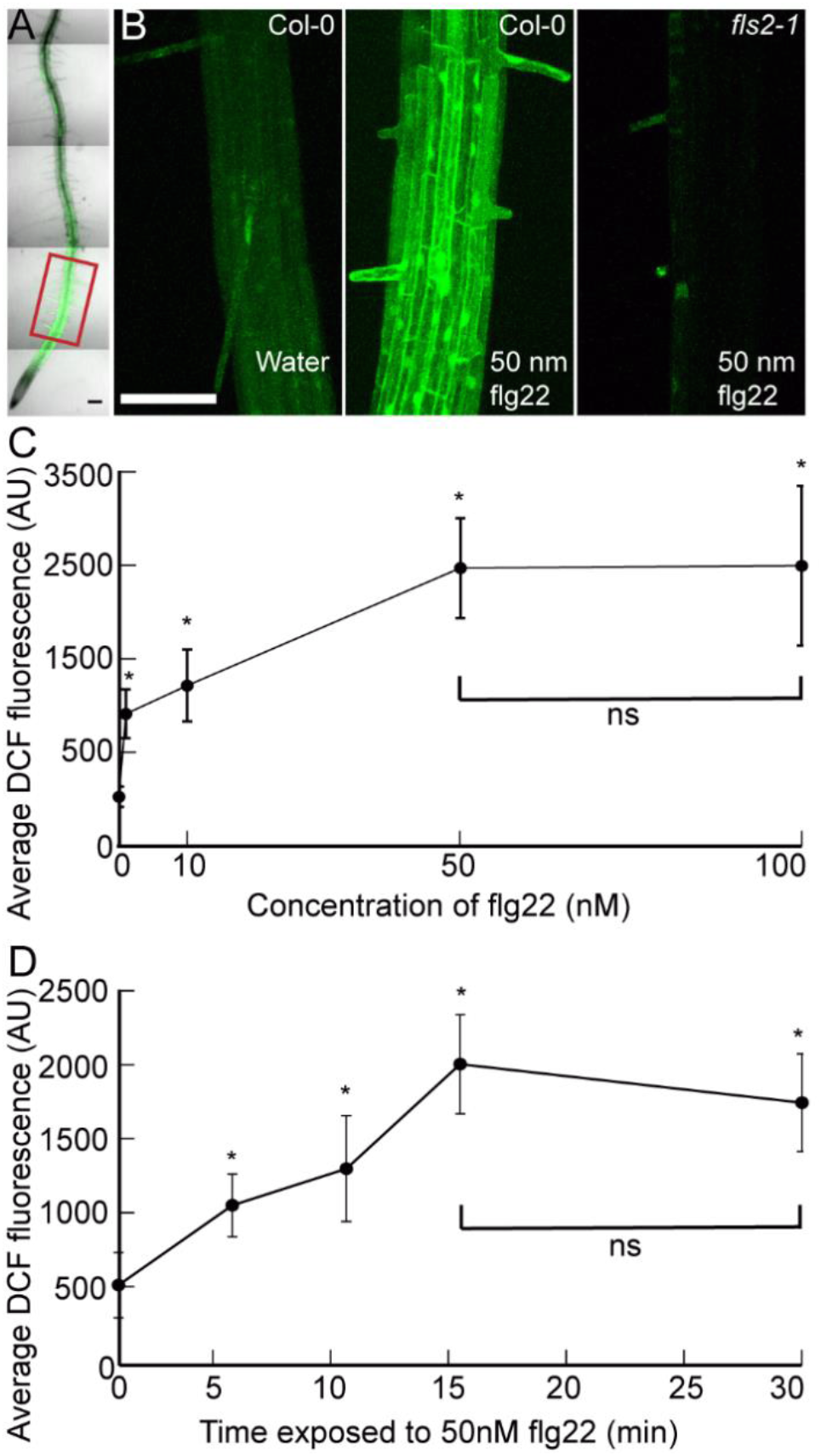
Dose and time effects of flg22 on ROS bursts in primary roots. A. Confocal micrograph showing flg22-induced ROS bursts by DCF staining of *Arabidopsis* primary root. Red box indicates region of interest used to quantify fluorescence. B. Confocal micrographs of the elongation zone of primary roots stained with DCF +/-flg22. C. Quantification of DCF-stained root tips showing dose response to different concentrations of flg22. D. Quantification of DCF-stained root tips showing timing of flg22-induced ROS bursts. Error bars represents standard deviation. Asterisks indicate significant difference (P < 0.01) between water and flg22 treatment determined by a two-way ANOVA followed by Tukey’s Posthoc test. Scale bars = 100 µm

### Analysis of flg22-induced G protein-specific phosphoproteome

Having established the time course and dose response of flg22-induced ROS production in roots, we treated 12-day-old WT Columbia (Col-0) as well as mutant plants deficient in Gα, Gβ, and two out of the three Gγ subunits in Arabidopsis (Song *et al*, 2018a): *gpa1-4*, *agb1*-2, *agg1*-1, and *agg2*-1 quadruple mutant (designated *quad* or G protein mutant hereafter) seedlings with water (mock) or 50 nM flg22 and collected samples at 0, 3, and 15 minutes to ensure we captured the changes in the phosphoproteome as flg22 response reaches its temporal peak. We then quantified protein abundance and phosphorylation level by performing two-dimensional liquid chromatography-tandem mass spectrometry (2D-LC-MS/MS) on Tandem Mass Tag (TMTpro) labeled peptides (McAlister *et al*, 2012; Song *et al*, 2018b; Hogrebe *et al*, 2018; Clark *et al*, 2021). From these samples, we quantified the levels 9,319 proteins and 31,593 phosphorylation sites arising from 6,591 phosphoproteins (**Fig 2A** and **Appendix Datasets 1&2**).

**Figure 2.**
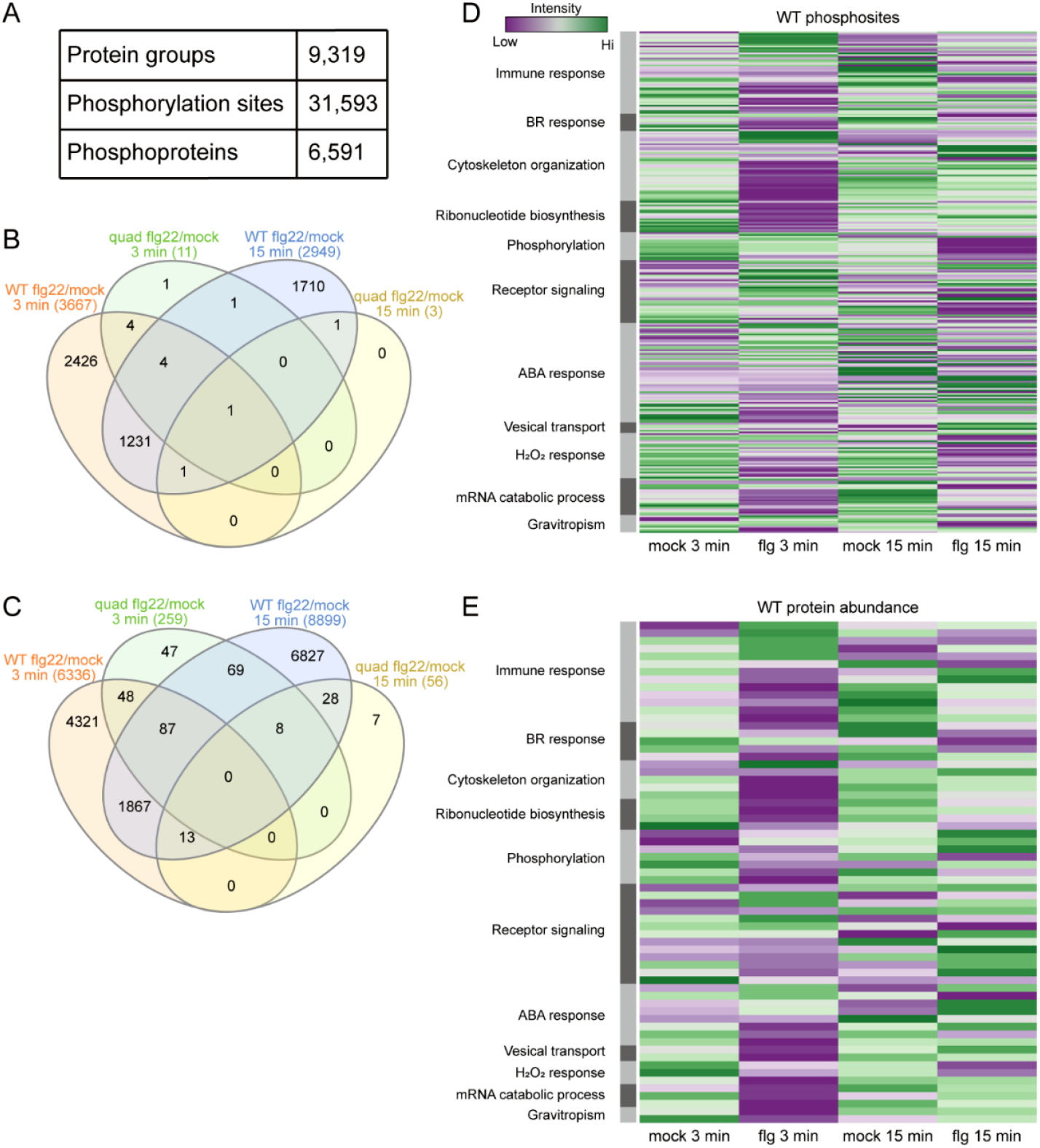
Analysis of a subset of At numbers for the dataset that appeared in previously published phosphoproteomes. A. The number of protein groups, phosphorylation sites, and phosphoproteins detected in this experiment B. Overlap between differentially abundant (q<0.1) proteins in WT and quad mutant backgrounds after 3 or 15 minutes of flg22 treatment. C. Overlap between differentially abundant (q<0.1) phosphorylation sites in WT and quad mutant backgrounds after 3 or 15 minutes of flg22 treatment. D. Comparing phosphorylation level of phosphosites in our phosphoproteomics dataset among selected 185 proteins in WT plants treated with mock (water) or flg22 for 3 or 15 minutes. Proteins were categorized using groups created by Gene Ontology enrichment analyses. E. Comparing protein abundance in our proteomics dataset among the 185 selected proteins in WT plants with mock (water) or flg22 for 3 or 15 minutes. Proteins were categorized using groups created by Gene Ontology enrichment analyses.

In WT plants, 6,336 sites corresponding to 2,278 phosphoproteins and 8,899 sites corresponding to 2,936 phosphoproteins were differentially phosphorylated following 3 minutes and 15 minutes, respectively, of flg22 treatment compared to the paired mock (H2O) samples. In the absence of the heterotrimeric G protein components (in the *quad* mutant), only 259 and 56 sites (corresponding to 104 and 29 phosphoproteins) are differentially phosphorylated after 3 and 15 minutes of treatment, respectively. These phosphosites represent only 3.1% (3 min) and 0.4% (15 min) of the flg22-regulated phosphoproteome of Col-0, suggesting the majority of the flg22-regulated sites are G-protein dependent (**Fig 2C, Appendix Dataset 2, and** Appendix Fig S1B). The results for protein abundance were similar, with a large number of protein groups differentially expressed after 3 minutes and 15 minutes of flg22 treatment in Col-0 (3,667 and 2,949, respectively) but minimal changes in protein abundance in the *quad* mutant (11 and 3, respectively) (**Fig 2B, Appendix Dataset 1, and Fig S1A**). These results indicate that the vast majority of flg22-induced proteome remodeling requires a functional G protein complex. The criteria for selection of 185 of these proteins is described in the Discussion. The changes in the phosphorylation (**Fig. 2D**) and abundance (**Fig 2E**) were analyzed for predicted function and are illustrated by heatmaps.

### Pathogenic effectors preferential target phosphoproteins

Highly connected nodes or hubs are preferential targets of pathogen effectors (Arabidopsis Interactome Mapping Consortium, 2011b; Mukhtar *et al*, 2011b; Wessling *et al*, 2014; González-Fuente *et al*, 2020), thus we performed network topology analysis of these flg22-induced phosphoproteome in the context of Arabidopsis protein-protein interactions. As described in the Methods section, we used an expanded version of the experimental interactome (Szklarczyk *et al*, 2015), and integrated the 3,734 flg22-induced phosphoproteins. The resulted network encompassed 6,618 nodes with 15,781 interactions including interactions from the Arabidopsis immune network and G-protein interactome (**Fig 3A**, **Appendix Dataset S3**). Network centrality analysis revealed that phosphorylated nodes possess high degree compared to non-phosphorylated proteins (**Fig 3B; Appendix Dataset S3**, Student’s *t*-test p-value ≤0.001). In other words, phosphorylated proteins have more interacting partners than non-phosphorylated proteins during flg22-induced defense response. This analysis inferred that the in-planta immune network and G-interactome utilize phosphorylation for efficacious downstream signaling during pathogen infection to modulate the host defense responses. Next, we computed the hubs (proteins with degree ≥ 15, designated hub_15_) in our flg22 induced protein-protein interaction experimental (PPIE). Hubs are enriched among phosphorylated nodes (**Fig 3C**, hypergeometric test p-value ≤ 0.001) i.e., the frequency of hub_15_ in phosphorylated nodes is statistically more prevalent than in non-phosphorylated nodes. Furthermore, we report that hub15 nodes are enriched in the immune network as a whole, which establishes our previous findings (Ahmed *et al*, 2018b). Concomitantly, we also observed that hub15s are statistically more prevalent in phosphorylated nodes of the G-interactome and immune network, whereas they are under-enriched (depleted) among the non-phosphorylated proteins during flg22 treatment (**Fig 3C**, **Appendix Dataset S3**, hypergeometric test p-value ≤ 0.001). Additionally, the functional analysis of phosphorylated hubs in immune and G-protein interactome nodes revealed that most of these proteins are significantly involved in stomata closure, defense response, freezing response, membrane docking, hormone signal transduction, and export from cell (**Fig 3D**, p-value≤0.05). Taken together, these network analyses confirmed the notion that flg22-dependent immune signal requires signal competent phosphorylated hubs.

**Figure 3.**
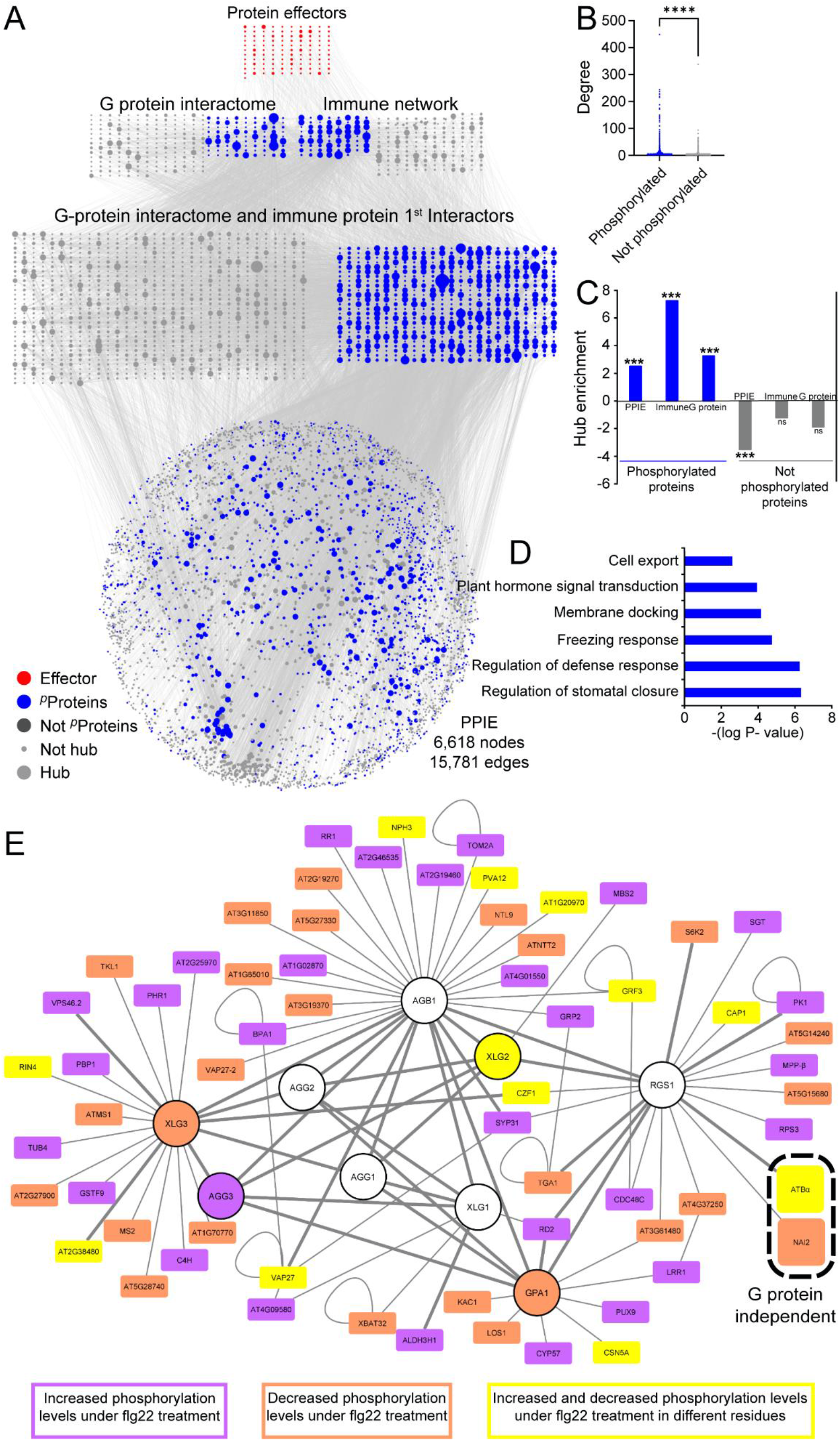
Pathogen-induced phosphoproteins are preferentially selected by effector proteins. A. The Arabidopsis experimental PPI (PPIE) network with 6,618 nodes and 15,781 interactions. (Red= effectors, blue= phosphorylated proteins (pProteins), Grey= not phosphorylated proteins (Not pProteins), large size= hub, small= not hub). B. The degree distribution of phosphorylated illustrates significantly high degree nodes than that of non-phosphorylated proteins in PPIE (Student *t*-test p-value ≤0.001). C. The significant hub enrichment of phosphorylated proteins and not phosphorylated proteins in PPIE, immune, and G protein interactome nodes (hypergeometric test enrichment p-value ≤ 0.001, ns = not significant). D. The functional enrichment of immune, and G protein interactome hub_15_ nodes (p-value≤ 0.05). E. Proteins from the Arabidopsis G Protein Interaction Database that respond to flg22 treat-ment. (mock vs. 3 or 15 min flg22, q < 0.1), having increased phosphorylation (purple), decreased phosphorylation (orange) or both (yellow) in different residues. The two high-lighted proteins represent the only proteins that respond to flg22 treatment on the quad mutant. Thicker edges represent a higher confidence of interaction based on previously published physical interactions. Arabidopsis G protein interactome can be accessed at http://bioinfolab.unl.edu/emlab/Gsignal.

### Arabidopsis G Protein Interactome contains proteins that respond to flg22

**Fig 3E** shows proteins in the Arabidopsis G-protein interactome that have flg22-induced changes in phosphorylation. After 3 minutes, flg22 treatment changes the abundance of phosphosites within 26 proteins that interact directly with heterotrimeric G protein components, including AtRGS1. This number is even higher with 15 minutes of treatment, when 43 proteins show significant differences in phosphorylation. Only two of those proteins show different phosphorylation patterns on the *quad* mutant and both of them are AtRGS1 interaction partners (**Fig 3E**, dashed oval outline) (Klopffleisch *et al*, 2011a). The PROTEIN PHOSPHATASE 2A (PP2A) regulatory B subunit (ATBα) is one of the two proteins that have decreased phosphorylation levels under stress in a G protein-independent manner. PP2A is a widely conserved serine/threonine phosphatase found in both the heterodimer and heterotrimer forms (Xu *et al*, 2006). The PP2A heterotrimer is composed of the structural A subunit, the substrate-recognition B subunit, and the catalytic C subunit. ATBα, the alpha isoform of the B subunit, along with ATBβ make up a subfamily of B subunits that are dissimilar to all other isoforms of the B subunit (Farkas *et al*, 2007). This finding raises the possibility that ATBα is one of the phosphatases in the G protein interactome that act antagonistically to the kinases that phosphorylate AtRGS1 in the presence of a signal (Tunc-Ozdemir *et al*, 2016). Regarding the heterotrimer, the alpha subunit AtGPA1 showed decreased levels of phosphorylation on a plant-unique serine residue (Ser202) over time and the gamma subunit AGG3 showed increased phosphorylated at Ser37 after 15 minutes of treatment compared to 3 minutes. flg22-induced post-translational modifications on G-proteins were found on non-canonical extra-large G proteins (XLGs), where XLG2 showed a decreased phosphorylation level on several residues outside the Gα domain (Ser75, Ser184, Ser185, Ser190, Ser191, Ser194 and Ser198) and an increase phosphorylation on residues Ser13 and Ser38 after 15 minutes. pSer13 and pSer38 are novel flg22-induced phosphorylation sites in roots, while Ser141, Ser148, Ser150 and Ser151 were previously detected as BIK1 phosphorylation sites under flg22 treatment in Arabidopsis protoplasts (Liang *et al*, 2016). This signaling dependency of the canonical G proteins, together with the XLG major modifications, prompts the hypothesis of regulation by competition between the XLGs and AtGPA1 for the AGB1/AGG dimers (Urano *et al*, 2016).

### Biochemical interaction between ATBα phosphatase and AtRGS1

Because we have no information about AtRGS1 phosphatases in the AtRGS1 phosphorylation-dephosphorylation cycle, we chose to examine more closely ATBα, a substrate recognition subunit of PROTEIN PHOSPHATASE 2A (PP2A). Its interaction with AtRGS1 *in planta* was tested with the Firefly Split Luciferase assay. As a negative control, we tested the interaction of AtGPA1 with AGB1 lacking its obligate Gγ subunit and observe no interaction (**Fig 4A**). As a positive control we used the interaction of the G protein heterotrimer subunits. AtRGS1 interacted with ATBα but not the ATBβ isoform suggesting that the ATBα interaction with AtRGS1 is isoform specific. Therefore, to quantitate and address specificity of the interaction of this substrate recognition, we tested *in vitro* interaction and phosphorylation of AtRGS1. We chose the well-characterized phosphorylation reaction by its kinase WNK8 (Urano *et al*, 2012b). In the absence of ATBα, WNK8 bound to and phosphorylated AtRGS1 as previously reported. However, the addition of the ATBα substrate-recognition subunit of this phosphatase blocked phosphorylation of AtRGS1 as expected for a substrate recognition subunit (**Fig 4B**). One interpretation of this observation is that ATBα is binding to its substrate and blocking phosphorylation. In this interpretation, we posit that ATBα is a phosphatase involved in the AtRGS1 phosphorylation-dephosphorylation cycle of the Ser/Thr phosphocluster necessary for endocytosis. The reciprocal experiment of showing that the PP2A-ATBα phosphatase complex dephosphorylates AtRGS1 is hampered by the heterotrimeric property of PP2A making it difficult to reconstitute a functional phosphatase *in vitro* (Farkas *et al*, 2007).

**Figure 4:**
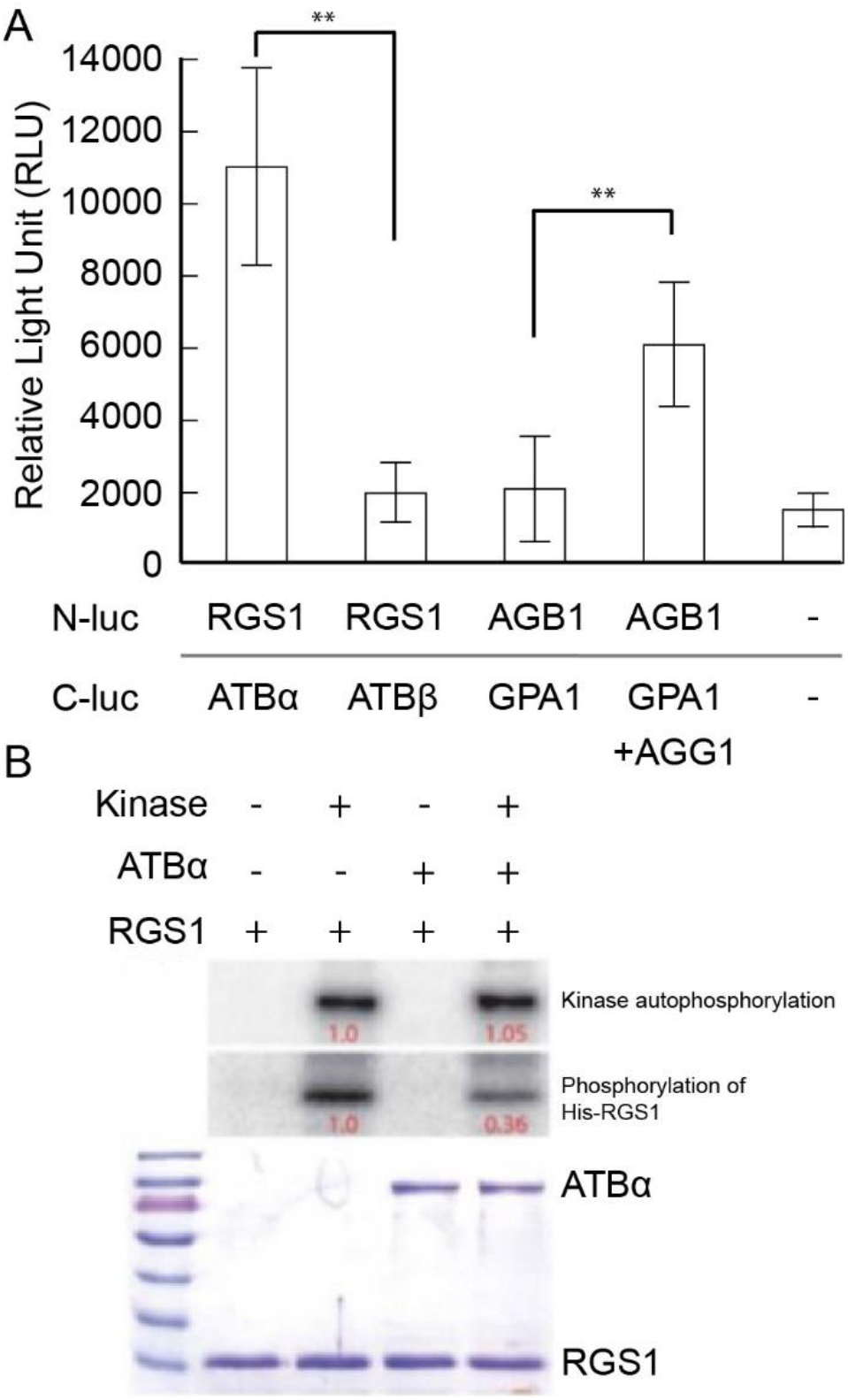
ATBα but not ATBβ interact with RGS1. A. Split luciferase assay showing protein interactions in vivo. ATBα but not ATBβ interacts with RGS1. Positive control is complementation by the heterotrimeric G protein complex (AtGPA1/AGB1/AGG1). Negative control is AtGPA1 and AGB1 in the absence of AGG1 as well as empty C- and N-luc vectors. Graphs are representatives of three experimental replicates. Error bars represent confidence intervals (CI). Asterisks indicate significant difference (P < 0.01) determined by a two-way ANOVA followed by Tukey’s Posthoc test. n = 36. B. ATBα decreases AtRGS1 phosphorylation catalyzed by a WNK-family kinase. 0.5 ug of GST-WNK8, 5 ug of GST-ATBα and/or 10ug of His-RGS1 were incubated in a 15ul of kinase reaction buffer (5mM Tris-HCl pH 7.5, 1 mM MgCl2, 0.4 mM ATP, 1mM PMSF) with a radio-labeled [γ-^32^P]-ATP at 20°C for 4hr. The samples were separated on a SDS-PAGE gel and exposed on a PHOSPHOR screen. The PHOSPHOR screen image and the Coomassie-stained gel are shown. The relative amounts of 32P were quantitated and provided as relative values in this figure.

### Genetic interaction between AtRGS1 and the ATBα phosphatase

Having shown *in vivo* and *in vitro* biochemical interaction between ATBα and AtRGS1, we tested genetic interactions. A quantifiable property of AtRGS1 is internalization by signals, including flg22 (Watkins *et al*, 2021). This is a quantifiable event that serves as a proxy for G protein activation (Fu *et al*, 2014a). Phosphorylation at the phosphocluster site leads to endocytosis, and thus if the phosphatase responsible for dephosphorylation of these phosphoserines is genetically ablated, we expect that the level of internalization is higher than expected even in the absence of flg22. Therefore, we crossed AtRGS1-YFP into the *atbα-1* loss-of-function mutant and quantitated flg22-induced endocytosis of AtRGS1 (Heidari *et al*, 2013). We included in our survey three other phosphatases shown to interact with AtRGS1 in a yeast two-hybrid screen (Klopffleisch *et al*, 2011b). These are ABSCISIC ACID INSENSITIVE 2 (ABI2), TYPE ONE PROTEIN PHOSPHATASE 8 (TOPP8), and DUAL SPECIFICITY PHOSPHATASE 1 (DSP1). ABI2 and TOPP8 are protein phosphatases that cause dephosphorylation events at serine and threonine residues (Farkas *et al*, 2007). ABI2 has been shown to negatively regulate abscisic acid (ABA) signaling in response to increased ABA by removing phosphate groups (Merlot *et al*, 2001). TOPP8 is an isozyme in the type one protein phosphatase family, many of which are predicted to act in cell cycle regulation (Farkas *et al*, 2007). Conversely, DSP1 is one of many dual-specificity phosphatases part of the protein tyrosine phosphatase (PTP) family which display varied enzymatic characteristics (Romá-Mateo *et al*, 2011).

As shown in **Fig 5A**, the basal level of AtRGS1 internalization at time zero was, as previously reported, about 20 % (Watkins *et al*, 2021) and was observed for wild type and *abi2-1*, *dsp1-4,* and *topp8-4* mutants cells (**Appendix Figure S2C**). In the absence of ATBα, the basal level of AtRGS1 was twice that for the other genotypes (P < 0.01) and nearly the maximum level of flg22- induced AtRGS1 endocytosis. Likewise, Ser/Tr phosphatase inhibitors, calyculin A (Stubbs *et al*, 2001) and cantharidin (Bajsa *et al*, 2015), accelerated RGS1-YFP endocytosis in the absence and presence of flg22 (**Appendix Figure S2A**). In contrast, RGS1-YFP internalization was not affected by treatment with the tyrosine phosphatase inhibitor, sodium orthovanadate (Yemets *et al*, 2008). Endocytosis was also blocked in the RGS13SA-YFP (**Appendix Figure S2B**), which contains 3 point mutations of Ser residues in the phosphocluster required for AtRGS1 endocytosis: S428A, S435A, and S436A (Urano *et al*, 2012a). This is consistent with the necessity of phosphorylated Ser/Thr residues to induce endocytosis (Urano *et al*, 2012a), and provides further evidence for the existence of a Ser/Thr phosphatase that dephosphorylates these residues. Taken together, these observations are consistent with the notion that ATBα is part of a phosphatase that is involved in the phosphorylation state of the phosphoserine cluster on AtRGS1.

**Figure 5:**
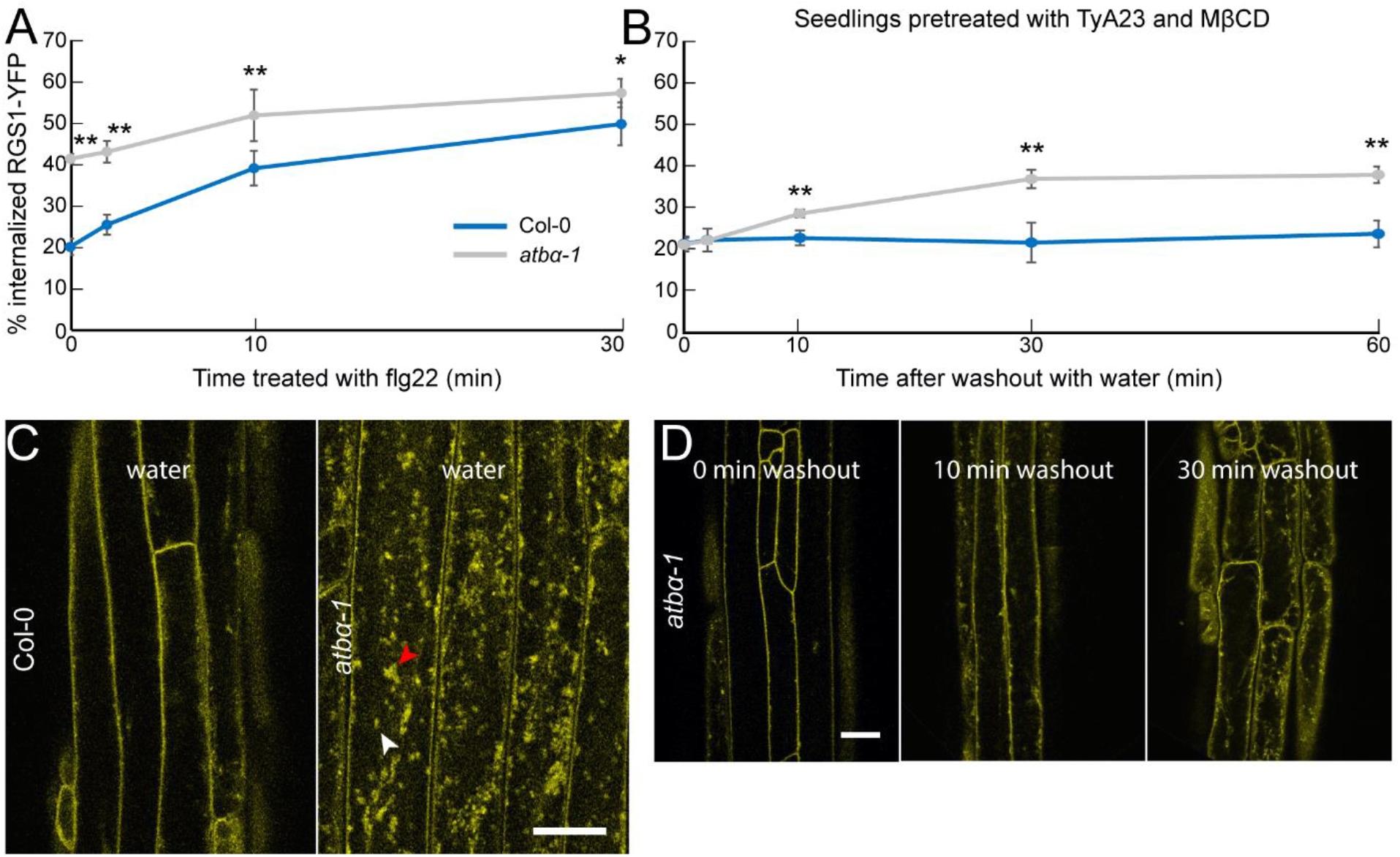
ATBα modulates RGS1 internalization in etiolated hypocotyl cells. A. flg22-induced RGS1-YFP internalization measured over time in Col-0 and *atbα-1* null mutant. ** and * represents statistical significance (P < 0.01 or P < 0.05 respectively) between Col-0 and *atbα-1* within timepoint. Error bars represent SEM. n=30 across three separate experimental replicates. B. Percent internalized RGS1-YFP measured in Col-0 and *atbα-1* 0, 2, 10, 30, or 60 minutes after washout with water. Col-0 seedlings were incubated in water for 2 hours and *atbα-1* seedlings were incubated for 2 hours with 50 µM TyrA23 and 5 mM MβCD. ** and * represents statistical significance (P < 0.01 or P < 0.05 respectively) between Col-0 and *atbα-1* within timepoint. Error bars represent SEM. n=32-48 across three separate experimental replicates. C. Representative confocal micrographs of *atbα-1* quantified in panel B. Scale bar=20 µm. D. Confocal micrographs of *atbα-1* expressing RGS1-YFP. Scale bar=20 µm.

To further compare the internalization dynamics of AtRGS1-YFP in the presence and absence of ATBα, we utilized endocytosis inhibitors to recapitulate the Col-0 phenotype in *atbα-1*. The two major endocytic pathways in plants, clathrin-mediated endocytosis (CME) and sterol-dependent endocytosis (SDE), are associated with AtRGS1-YFP internalization and activation of downstream targets of G-protein signaling (Watkins *et al*, 2021), including ROS bursts (Li *et al*, 2012; Dhonukshe *et al*, 2007). To decrease the basal level of internalized AtRGS1-YFP in *atbα-1*, we incubated seedlings for two hours with 50 µM Tyrphostin A23 (TyrA23) and 5 mM methyl-β-cyclodextrin (MβCD), which inhibit CME and SDE, respectively (Ohtani *et al*, 1989; Ilangumaran & Hoessli, 1998; Banbury *et al*, 2003; Dhonukshe *et al*, 2007; Watkins *et al*, 2021), resulting in basal level of AtRGS1 internalization similar to Col-0 (**Fig 5B, C**). This finding is consistent with AtRGS1-YFP phosphorylation driving endocytosis via CME and SDE pathways (Watkins *et al*, 2021). After washing out TyrA23 and MβCD with water, we observed AtRGS1-YFP internalization over a 60-minute time course resulting in a return to ∼40% internalized protein, the *atbα-1* basal state of internalization (**Fig 5B, C**). Additionally, the special pattern of basal AtRGS1-YFP internalization in *atbα-1* is similar to Col-0 treated with flg22 or D-glucose as previously reported (Watkins *et al*, 2021), with *atbα-1* containing both endosome-(**Fig 5C**, white arrows) and micro-domaine-like (**Fig 5C**, red arrows) internalized structures. D-glucose-induced internalization was 20% higher in *atbα-1* compared to WT (P<0.01) (**Appendix Figure S2D**).

### Loss of ATBα results in AtRGS1-YFP degradation

Because AtRGS1 phosphorylation is required for internalization and subsequent degradation, the absence of phosphatase activity on this protein is expected to promote lower AtRGS1 levels within the cell. Cycloheximide was used to inhibit protein synthesis in Arabidopsis seedlings treated with flg22 for 1 and 2 hours. Confocal microscopy revealed a 70% reduction in AtRGS1-YFP fluorescence levels in *atbα-1* compared to WT (**Fig 6A**). Additionally, the RGS1-YFP reporter in WT plants showed increased protein signal when compared to plants crossed with the *atbα-1* allele, indicating that AtRGS1 levels are naturally lower in the absence of the phosphates subunit that is responsible for the negative regulation of the internalization process (**Fig 6A, B**). Pretreatment with the endocytosis inhibitors, TyrA23 and MβCD, recapitulated the WT phenotype in *atbα-1* treated with cycloheximide. This suggests that AtRGS1-YFP internalization results in degradation of the protein, and the ATBα negatively regulates this process. To further validate this finding, we analyzed AtRGS1-YFP abundance in protein extracts from whole plants via immunoblot analysis (**Fig 6C, D**). We found that cycloheximide treatment decreased AtRGS1-YFP protein abundance in *atbα-1* by nearly 100%. The discrepancy of 70% reduction (by fluorescence quantitation) vs. 100% (by western quantitation) is likely due to the greater dynamic range of confocal microscopy compared to immunoblot quantitation. Nonetheless, in both cases, we observed a vastly reduced steady-state AtRGS1-YFP protein level in *atbα-1*.

**Figure 6:**
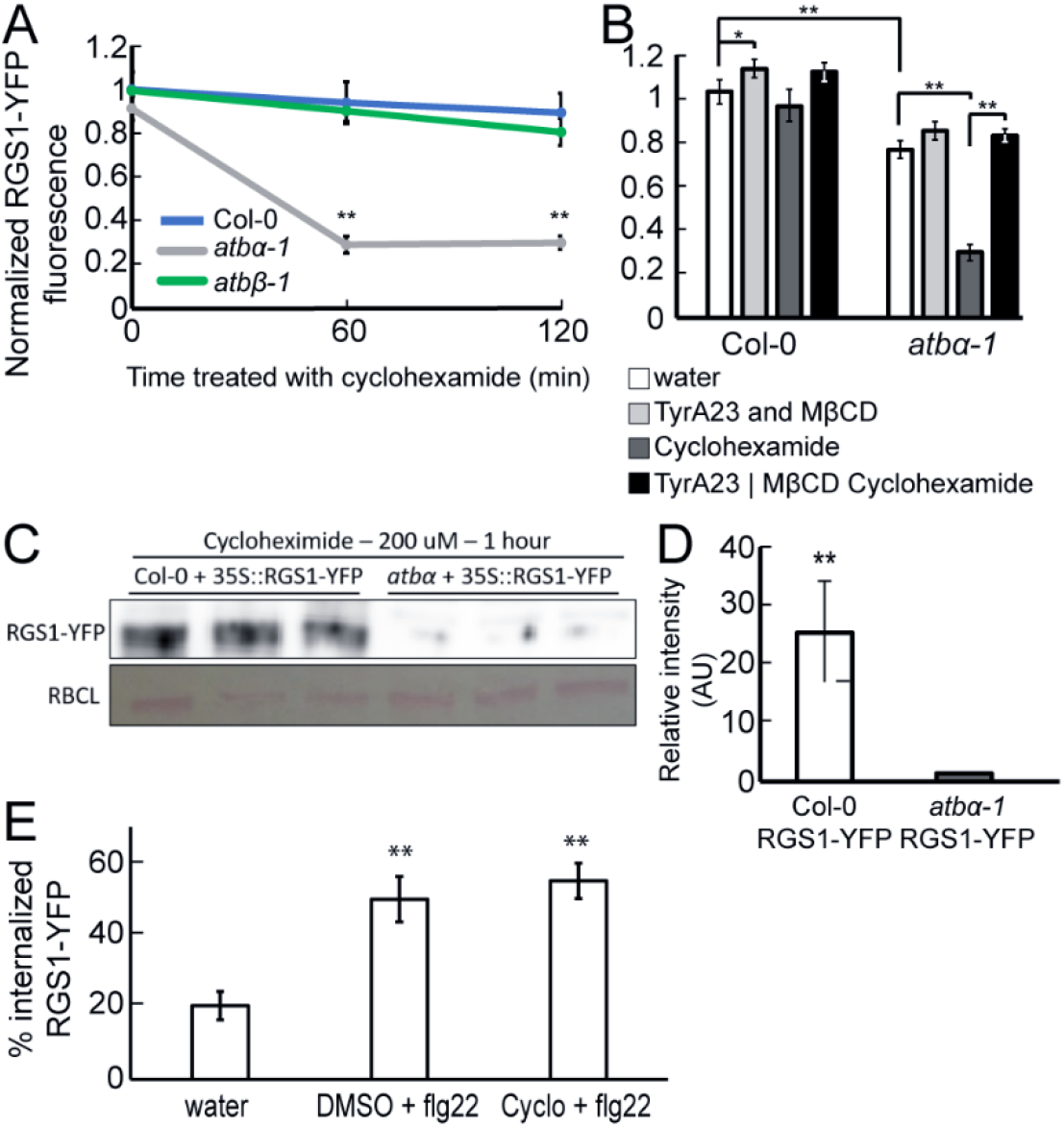
flg22-induced internalization of RGS1 leads to degradation. A. RGS1-YFP protein abundance measured by YFP fluorescence in Col-0, *atbα-1*, and *atbβ-1* treated with 200 µM cycloheximide for 0, 60 and 120 minutes. ** represents statistical significance (P < 0.01) between Col-0 and *atbα-1* within timepoint. Error bars represent CI. n=25-49 across three separate experimental replicates. B. RGS1-YFP protein abundance measured by YFP fluorescence in Col-0 and *atbα-1* after treatment with inhibitors. Seedlings were either treated with water, TyrA23 and MβCD, or cycloheximide for 1 hour, or they were pretreated with TyrA23 and MβCD for one hour followed by cycloheximide treatment for 1 hour prior to imaging. ** and * represents statistical significance (P < 0.01 or P < 0.05 respectively). n=25-49 across three separate experimental replicates. C. Western blots of RGS1-YFP protein extracts from whole seedlings of Col-0 and *atbα-1* after treatment with 200 µM cycloheximide for 60 minutes. D. Western blot quantification of AtRGS1-YFP normalized by RuBisCO levels. **\*\*** represents statistical significance (P<0.01) compared to control (RGS1-YFP/Col-0) and determined by unpaired t-test. E) RGS1-YFP internalization in response to 30 minutes of flg22 treatment after pretreatment with DMSO or 200 µM cycloheximide for 60 minutes. ** represents statistical significance (P < 0.01) between water and treatment. Error bars represent CI. n=25.

Finally, the role of tonic cycling in modulating the percent internalized AtRGS1-YFP in response to flg22 was assessed. To accomplish this, flg22-induced AtRGS1-YFP endocytosis in the presence and absence of cycloheximide was compared. AtRGS1-YFP endocytosis levels were 2.5 and 2.75 times higher in flg22 treated seedlings with and without cycloheximide, respectively; however, we did not find a significant difference when comparing flg22-treated seedlings in the presence or absence of cycloheximide (**Fig 6E**). This suggests that nascent AtRGS1-YFP is not substantially transported to the membrane during the 30-minute flg22 treatment. This phenomenon could be one mechanism the cell uses to dampen itself to recurring G protein-coupled signals.

### ATBα modulates plant immune response and development

Given that ATBα interacts with and modulates the phosphorylation and subsequent internalization of AtRGS1, we hypothesized that such functions would affect the activation of downstream G protein signaling targets. One of the most rapid known events in flg22-dependent G protein signaling is the ROS burst, beginning within seconds of recognition of the MAMP, and peaking between 10 and 15 minutes before returning to the base line within 60 minutes. Because of the involvement of G protein complexes in flg22-induced ROS production (Ishikawa, 2009; Lorek *et al*, 2013), we quantitated flg22-induced ROS production in the *atbα-1* null mutant and compared to *rgs1-2* and Col-0. The peak of ROS production induced by 100 nM flg22 treatment was enhanced in *rgs1-2* compared to WT (**Fig 7A**), consistent with previous reports (Ghusinga *et al*, 2021b). *atbα-1* mutant also showed an enhanced ROS peak compared to WT consistent with increased RGS1 internalization and degradation observed in the mutant.

**Figure 7.**
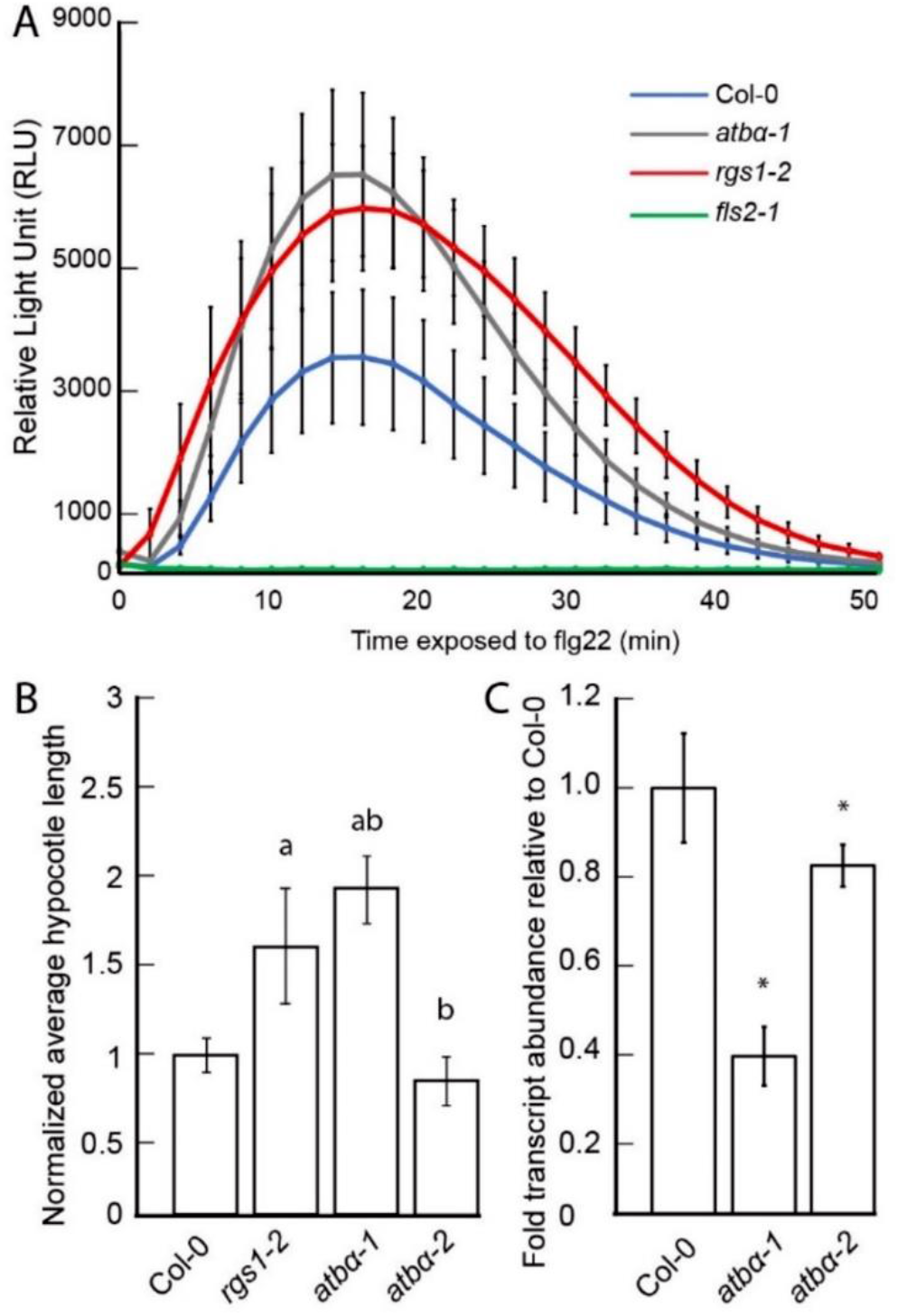
ATBα modulates plant immune response and development. A. flg22-induced ROS, reported as Relative Luminescence Units (RLU), in leaf disks generated from 5-week-old plants treated with 100 nM flg22. Error bars represent CI. Graph is representative of three separate experiments. n = 20–35. B. Hypocotyl lengths of Col-0, *rgs1-2*, *atbα-1*, and *atbα-2*. (a) denotes significantly difference from Col-0 (P<0.01). (b) denotes significant difference between phosphatase mutant alleles and *rgs1-2* (p<0.01) determined by a two-way ANOVA followed by Tukey’s Posthoc test. n = 35-50. CRT-qPCR analysis of Col-0, *rgs1-2*, *atbα-1*, and *atbα-2*. Averages of three biological replicates are reported. ** represents significant difference (P < 0.01) between Col-0 and mutants as assessed by ANOVA and Tukey’s posthoc test.

To further characterize differences in the phosphatase mutants related to AtRGS1-dependent G-protein signaling, we measured hypocotyl lengths in etiolated seedlings, a G protein-dependent phenotype (Chen *et al*, 2003), and compared them to *rgs1-2*. Etiolated *rgs1-2* seedlings have elongated hypocotyls compared to WT and this is associated with upregulated G protein activity (Chen *et al*, 2003). Therefore, if one of these phosphatases modulates G protein signaling, then we would expect changes in hypocotyl growth. These established phenotypes were used to reveal potential regulatory effects of phosphatases based on their respective mutant phenotype. Generally, the phosphatase mutant alleles resembled the hypocotyl lengths of *rgs1-2*, which were significantly longer than Col-0 (**Fig 7B**). The one exception was *atbα-2*, an intron insertion allele, which showed a WT phenotype. Transcriptional analysis showed that all phosphatase mutant alleles showed lower transcript levels than WT, but *atbα-2*, an intron insertion allele, had transcript levels 83% of WT (**Fig 7C**). The correlation between gene expression of the phosphatase mutant alleles and the hypocotyl elongation phenotype confirm that the phenotype is conferred by loss of ATB*α*.

## Discussion

From the vantage of phosphorylation states, innate immunity in roots is shown here to be rapid responding and dynamic. While it has been known since the beginning of plant G protein research that G signaling played some role in innate immunity, our systems analysis makes it clear how profound is that role. Many new avenues to dissect innate immunity are revealed. One of these is a phosphatase that regulates the steady-state phosphorylation status of a key modulator of G signaling. Because the action of this phosphatase leads to a change in G signaling, the second level of dynamics in pattern-triggered immunity was peeled away. Specifically, as AtRGS1 levels change with a time constant of minutes, the phosphorylation-dephosphorylation cycle occurring in seconds is predicted to change.

The foundation experiments designed to capture the relevant window of time and physiological concentration of flg22 for our subsequent phosphosite mapping also reveal new information about innate immunity in the root. The spatial and temporal information for flg22-induced signaling was achieved using the fluorescent ROS sensor, DCF (Halliwell & Whiteman, 2004a; Tyburski *et al*, 2009; Chapman & Muday, 2021). Confocal micrographs showed peak fluorescence in the elongation zone (EZ) (**Fig 1A**), which is consistent with a previous study that demonstrated that the EZ is especially sensitive to flg22 (Millet *et al*, 2010). Additionally, the EZ, but not the root tip, exhibits high gene expression of *FLS2*, the gene encoding the canonical flg22 receptor (Beck *et al*, 2014). This is consistent with our results where we did not see significant increases in flg22-induced ROS in root tips, further suggesting that this region is largely insensitive to flg22. Taken together, we chose to image flg22-induced ROS in the EZ to establish the dose and treatment time of flg22 for phosphoproteome analysis.

No significant change in AtRGS1 phosphorylation was flagged as significant at the stringency set here which does not support considerable phosphonull mutant data suggesting that phosphorylation of Serine 431 is induced by flg22 treatment in Arabidopsis protoplasts (Liang *et al*, 2018). This may be due to the lack of resolution at this site due to the many validated phosphoresidues that are densely packed in this region of the C-terminal tail of RGS1 designated the phosphocluster, thus resulting in poor localization scores in our dataset. Also, the low abundance of AtRGS1 within the plant cell, together with the fast turn-over of AtRGS1, may limit significant differences in AtRGS1 phosphorylation using phosphoproteomics approaches (Tunc-Ozdemir *et al*, 2017). However, we do not discount other possibilities such as different flg22 concentrations, time of the treatments, and tissue.

The major role of the heterotrimeric G protein is even clearer in the dataset analysis of well-known components of PAMP recognition and signaling in plants (Tunc-Ozdemir & Jones, 2017). After flg22 recognition by the LRR-RLK, FLS2, an active complex forms with another LRR-RLK, BAK1, which then phosphorylates the receptor-like cytoplasmic kinase BIK1 (Chinchilla *et al*, 2007). Once phosphorylated, BIK1 triggers a ROS burst response and activates MAPK cascades (Lu *et al*, 2010). This dataset shows that the expected differentially increased levels of phosphorylation from BIK1 and also from the two threonines and tyrosines from each MPK3 and MPK6 (Rayapuram *et al*, 2018) only occur on the wild type plants and not on the quad mutant, consistent with the notion that major G protein components act upstream of characterized components of plant immunity. The time course profiles also revealed an additional set of receptors in innate immunity. Among those kinases in **Appendix Dataset S4**, some are expected such as WALL-ASSOCIATED KINASE (Faris & Friesen, 2020), NSP-INTERACTING KINASE 1 (NIK1, (Teixeira *et al*, 2019; Gouveia *et al*, 2017), CHITIN ELICITOR RECEPTOR KINASE 1 (Desaki *et al*, 2018), BAK1 RELATED KINASE (Tunc-Ozdemir & Jones, 2017), FERONIA (FER,(Yang *et al*, 2020), and HAESA (Patharkar *et al*, 2017), while 22 others present opportunity. Our results were refined with data from the Arabidopsis experimental protein-protein interaction network, the Arabidopsis Immune Network, and the Arabidopsis G Protein Interactome database (AGIdb). The AGIdb was generated in a previous study by investigating interactions among proteins involved in the G protein pathway, using a yeast-two-hybrid approach combined with bimolecular fluorescence complementation (Klopffleisch *et al*, 2011b). This interactome contains 4 ser/thr phosphatases that interact with AtRGS1, including ATBα, ABI2, TOPP8, and DSP1. Of the four phosphatases, only ATBα showed differential phosphorylation in response to flg22 treatment with some residues showing increased phosphorylation levels while other residues showed reduced phosphorylation levels (**Fig 3G**). These changes were present in WT and the *quad* mutant, suggesting they are G protein independent.

We used three published flg22-induced datasets that each utilized different treatment times, tissues. We also compared these results with a xylanase, and oligo-galacturonide (pectin fragments) induced phosphoproteome, respectively (**Appendix Table S1**) (Benschop *et al*, 2007; Nühse *et al*, 2007; Rayapuram *et al*, 2014; Kohorn *et al*, 2016). The analysis required that a phosphoprotein to contain differentially expressed phosphosites(s) in at least one other published dataset. This resulted in 185 phosphoproteins (**Appendix Dataset S5**) that were present in two or more datasets despite major differences in experimental design and lesser proteome coverage of the earlier technology. Heatmaps were generated to visualize changes among these 185 phosphoproteins in response to flg22 treatment in WT (**Fig 2D and E**). Consistent with the quick ROS burst induced by flg22, gene ontology (GO) terms known to be impacted by flg22 signaling were enriched. For example, flg22 decreased the phosphorylation of kinases and cytoskeleton proteins. Other groups, like ABA response and immune response, contain sets of proteins whose phosphorylation increases in response to flg22, and others in which it decreases in response to flg22. There are also groups that are time point dependent. For example, vesicle transport phosphorylation decreases at 3 minutes, but increases at 15 minutes. Gravitropism also seems to have an opposite response between time points.

In particular, one study focused on changes in phosphorylation in response to OGs (Kohorn *et al*, 2016). A 5-minute treatment was sufficient to induce phosphorylation of 50 different proteins and, of this set, ATBα was phosphorylated at two serine residues. Comparing the OG results to our dataset, the first of these residues was phosphorylated after 15 minutes of flg22 in both WT and the *quad* mutant, suggesting an overlapping role of ATBα in the OG and flg22 response pathways. Additionally, the second residue was phosphorylated after 5 minutes of flg22 treatment in WT, but not detected in the absence of the G protein complex.

Among the genes present in numerous datasets, the protein encoded by *OPEN STOMATA2 (OST2)* was found to be phosphorylated in response to flg22 in two datasets as oligo-galacturonide concentrations (pectin fragments). OST2 is integral to induce stomatal closure in response to ABA (Merlot *et al*, 2007), which is important for plant growth in response to drought as well as bacterial invasion. Our results suggest the OST2 is jointly regulated via ABA and flg22 to illicit stomatal closure via two different stimuli. Another protein of interest across studies is FER. FER phosphorylation was altered in 3 out of 4 flg22 phosphoproteomes. In guard cells, FER interacts directly with the Gβγ dimer of the heterotrimeric G protein complex (Yu *et al*, 2018). Other notable genes found in overlapping phosphoproteomes are genes related to MAPK and Ca^2+^ signaling, including MAPK4 and 6 and CPK5, 9, and 13.

Given that plant pathogen effectors target the highly connected hub proteins preferentially than other proteins of interactome (Mukhtar *et al*, 2011b; Mishra *et al*, 2017b; Ahmed *et al*, 2018b; González-Fuente *et al*, 2020), we investigated the 3,734 phosphoproteins interactions in Arabidopsis experimental PPI (PPIE) network (Klopffleisch *et al*, 2011b; Szklarczyk *et al*, 2015; Mott *et al*, 2019; González-Fuente *et al*, 2020). The PPIE contains 257 G-proteins interactome nodes, 235 immune network nodes, 111 effector nodes and other proteins. Pathogen effectors target the highly connected nodes more efficiently to hijack the host system, the next line of investigation was to perform network topology analysis, specifically degree centrality to calculate the total interactions (Mishra *et al*, 2021b). We show that the phosphorylated proteins possess significantly high degree distribution than the non-phosphorylated proteins (**Fig 3B**; Student *t*-test p-value ≤0.001) suggesting the potential roles of highly connected nodes in efficacious signal transduction in response to pathogens or pathogen-mimic stimuli. As a result, we found 352 proteins as hub15 (nodes with ≥15 interactions) of which 278 are phosphorylated in our analysis. Further, we expanded our analysis of hub15 in Arabidopsis immune network and G-protein interactome nodes, and report that irrespective of group, the phosphorylated hubs are significantly over-enriched as compared to unphosphorylated hubs which are under-enriched in PPIE, immune and G-protein interactome nodes (**Fig 3C**, hypergeometric test p-value ≤ 0.001). Collectively, these results further advance the importance of G-proteins as highly connected signal transducer host proteins and potential pathogen targets in plant immune network. Overall, our network analysis also corroborates with the findings that majority of flg22-induced proteome relies on G-proteins.

As such, we compared phosphoproteins in our dataset to the experimental interactome and immune network (**Fig 3A**). This highlighted a set of targets of pathogen effectors, suggesting the involvement of these genes in the flg-22 induced phosphoproteome and the pathogen’s targeting of them to suppress pattern-triggered immunity. Among the connections between genes in our dataset and these two interactomes, we discovered the same interactions of G-proteins and effectors, including PP2AA2, a structural subunit of PP2A. PP2A is comprised of 3 structural subunits: A1, A2, and A3 (Farkas *et al*, 2007). We found that the substrate recognition subunit, A2, was phosphorylated after 15 minutes of flg22 treatment, and that phosphorylation levels were reduced in the *quad* mutant, suggesting these phosphosites are dependent on the G-protein. Interestingly, A2 is linked to plant defense pathways (Ahn *et al*, 2007). A2 also has a role in mitigating vesical trafficking via PIN proteins (Karampelias *et al*, 2016; Dai *et al*, 2012) and is involved in redox signaling in peroxisomes (Kataya *et al*, 2015). When considering the heterotrimeric nature of PP2A together with the diverse pathways that the A and B subunits of PP2A are associated with, it is possible that the PP2A phosphatase can regulate numerous parts of the flg22 pathway via different arrangements of subunits. Future studies should examine which structural and catalytic subunits interact with ATBα to regulate flg22-induced endocytosis.

Taken together, biochemical (**Fig 4**), bioinformatic (**Fig 3E**), physical (**Fig 4**), and genetic (**Fig 5, 7**) data place one of the dynamically phosphorylated targets, ATBα, in G-protein-dependent plant immunity, ends the long search for the AtRGS1 phosphatase, and congeals several observations in innate immunity. Arabidopsis encodes 17 substrate-specificity B subunits of PP2A, which interact in a variety of combinations to yield differentially-regulated outcomes (Jonassen et al., 2011; Trotta et al., 2011) and are broken down into 3 subfamilies. ATBα and ATBβ, make up their own subfamily characterized by 5 similar WD40 repeats that are unlike all other isoforms of the B subunit (Farkas *et al*, 2007). B subunits of PP2A are directly related to stress signaling (Trotta et al., 2011), and root growth (Blakeslee et al., 2008). PP2A is also a negative regulator of the PAMP-triggered immune response in Arabidopsis (Segonzac *et al*, 2014). Here, the PP2A holoenzyme composed of A1 and B’η/ζ was found to inhibit PAMP-triggered immune responses via association with BAK1, a key immune component of the flg22-signaling pathway (Sun *et al*, 2013; Toorn *et al*, 2012). Genetic ablation of some B subunits led to increased steady-state BAK1 phosphorylation and flg22-induced ROS bursts.

To our knowledge, PP2A (ATBα type) type is the first phosphatase associated with a 7-transmembrane GPCR-like protein. One other PP2A (B56δ type) was discovered by a phosphoproteomic screen to be activated by cAMP and may be important in GPCR-based signaling but it is not known if this phosphatase dephosphorylates a GPCR (Tsvetanova *et al*, 2021). Both B56δ and ATBα are B subunits as described above, however, ATBα has a 5-bladed WD40-repeat scaffold while B56δ has a B56 scaffold. Nonetheless, our study prompts a set of experiments for animal G signaling mechanisms.

In conclusion, our study provides a novel role for ATBα and the heterotrimeric PP2A phosphatase in regulating flg22-induced, G protein-dependent signaling. We also showed that the ATBβ isomer does not share this functionality despite being in the same subfamily with ATBα and containing a similar peptide sequence. Furthermore, our phosphoproteome dataset provides a detailed account of the G protein-dependent phosphorylation events that occur early in the flg22 signaling pathway.

## Methods

### Plant genotypes and growth conditions

Arabidopsis (*Arabidopsis thaliana*) ecotypes for all plants used in this study are Columbia-0 (Col-0). The *rgs1-2* (SALK_074376.55.00) protein-null, T-DNA-insertion mutant was created as previously described (Chen et al., 2003a). The *fls2-1* (SAIL_691_C4) protein-null, T-DNA-insertion mutant was created as previously described (Zipfel et al., 2004). The AtRGS1-YFP reporter was combined with *abi2-1* (SALK_015166C), *abi2-2* (SAIL_547_C10), *atbα-1* (SALK_032080C) (Heidari *et al*, 2013), *atbα-2* (SALK_090040C), *topp8-2* (SALK_125184), *topp8-4* (SALK_076144), *dsp1-3* (WiscDsLo473B10), *dsp1-4* (SAIL_116_C12), and *atbβ-1* (GK-290G04-01) mutant backgrounds through crossing to create stable transformants. Genotyping utilized primers provided in **Appendix Table S2**.

Unless otherwise described, seeds were surface sterilized with 70% ethanol for 5 minutes while vortexing followed by a 5-minute treatment with 95% ethanol. Seeds were subsequently washed 3X with dH20 ultrapure H20 and suspended in 12-well cell culture plates with ¼ MS with no sugar at pH 5.7 or plated in similar media with 0.8% agar added. Plates were wrapped in aluminum foil, cold-treated at 4°C for 2 days prior to germination.

For phosphoproteomics analysis, *Arabidopsis thaliana* accession Columbia (Col-0) was used as wild-type (WT) while the quadruple (quad) mutant consisted of *gpa1-4*, *agb1-2*, *agg1-1*, and *agg2-1* plants. After the surface sterilization seeds were placed on sterile nylon mesh (Amazon, Nylon 6/6 Woven Mesh Sheet, Opaque Off-White, 40" Width, 10 yards length, 110 microns mesh size# B0013HNZJC) that was on the top of Murashige and Skoog basal agar media with 0.5% sucrose and stratified at 4 C for 2 days in the dark. Plants were then grown for 12 days in a growth chamber with 24-hour constant light at the intensity of 150 photons per m^2^. After 12 days, 10ml of the mock (water) or flg22 solution was added directly on the roots and kept submerged for 3 min and 15 min. Root tissue was then harvested and flash frozen in liquid nitrogen. Prior to protein extraction the tissues were ground for 15 minutes under liquid nitrogen using a mortar and pestle.

### Chemicals

Methyl-β-cyclodextrin was purchased from Frontier Scientific and tyrphostin A23 (TyrA23) was purchased from Santa Cruz Biotechnology. All chemicals were indicated by the vendors to be >98% pure.

### Imaging ROS with Confocal microscopy

Chloromethyl 2’,7’-dichlorodihydrofluorescein diacetate, H2DCF-DA (Thermo Fisher), was used as a generic ROS sensor (Halliwell & Whiteman, 2004b). H2DCF-DA was prepared as previously described (Watkins *et al*, 2017, 2014). Briefly, H2DCF-DA dissolved in dimethyl sulfoxide to yield a 50 mM stock and diluted in deionized water to yield a final concentration of 6.25 µM. Roots were incubated with flg22 for the indicated time prior to a brief washout with water and a 10-minute incubation with the H2DCF-DA solution.

DCF fluorescence was imaged on a Zeiss 880 laser scanning confocal microscope and excited with 0.2% maximum laser power at 488 nm with a 2.0 digital gain and a Plan-Neofluar 20x/0.50 Ph2 objective lens. The DCF signal was collected between 495 and 550 nm with a pinhole yielding 1 Airy Unit, making sure to limit excess exposure to the laser that induces ROS. Maximum intensity projections (MIP) were produced from Z-stacks. All micrographs within each panel were acquired using identical offset, gain, and pinhole settings using the same detectors. DCF fluorescence intensities were measured in the MIPs using Fiji ImageJ by placing an ROI around the elongation zone near the root tip. The average intensity values within each ROI were recorded and averaged.

### Protein Extraction and Digestion

The proteomics experiments were carried out based on established methods (Song *et al*, 2018a; Clark *et al*, 2021). Protein was extracted and digested into peptides with trypsin and Lys-C using the phenol-FASP method as previously detailed (Song *et al*, 2018b, 2018b). Resulting peptides were desalted using 50 mg Sep-Pak C18 cartridges (Waters), dried using a vacuum centrifuge (Thermo), and resuspended in 0.1% formic acid. Peptide amount was quantified using the Pierce BCA Protein assay kit.

### Tandem Mass Tag (TMT) Labeling

The TMT labeling strategy used in this experiment is provided in **Appendix Dataset S6**. Approximately 40 µg of peptides were taken from each individual sample and then pooled. TMTpro™ 16plex labeling reagents (ThermoFisher, Lot UH290430) were used to label 150 µg of peptides, from each sample or pooled reference, at a TMT:peptide ratio of 0.2:1 as described in (Song et al., 2020). After 2 hours incubation at room temperature the labeling reaction was quenched with hydroxylamine. Next, the 16 samples were mixed together stored at -80°C until phosphopeptide enrichment. Labeling efficiency was checked by performing a 60-minute 1D run on 200 ng of TMT-labeled peptides. All samples had labeling efficiencies ≥ 97%.

### Phosphopeptide Enrichment

The TMT-labeled phosphopeptides were first enriched using the High-Select TiO_2_ Phosphopeptide Enrichment Kit (Thermo) using the manufacturers protocol. The High-Select Fe-NTA Phosphopeptide Enrichment Kit (Thermo) was then used on the flowthrough from the TiO_2_ enrichment to enrich additional phosphopeptides. The manufacturers protocol for the Fe-NTA kit was used except the final eluate was re-suspended with 50 μL 0.1% formic acid. The eluates from the TiO_2_ and Fe-NTA enrichments were combined and stored at -80°C until analysis by LC-MS/MS.

### LC-MS/MS

An Agilent 1260 quaternary HPLC was used to deliver a flow rate of ∼600 nL min-1 via a splitter. All columns were packed in house using a Next Advance pressure cell, and the nanospray tips were fabricated using a fused silica capillary that was pulled to a sharp tip using a laser puller (Sutter P-2000). 10 μg of TMT-labeled peptides (non-modified proteome), or ∼15 μg TiO_2_ or Fe-NTA enriched peptides (phosphoproteome), were loaded onto 10 cm capillary columns packed with 5 μM Zorbax SB-C18 (Agilent), which was connected using a zero dead volume 1 μm filter (Upchurch, M548) to a 5 cm long strong cation exchange (SCX) column packed with 5 μm Poly-Sulfoethyl (PolyLC). The SCX column was then connected to a 20 cm nanospray tip packed with 2.5 μM C18 (Waters). The 3 sections were joined and mounted on a Nanospray Flex ion source (Thermo) for on-line nested peptide elution. A new set of columns was used for every sample. Peptides were eluted from the loading column onto the SCX column using a 0 to 80% acetonitrile gradient over 60 minutes. Peptides were then fractionated from the SCX column using a series of 17 and 8 salt steps (ammonium acetate) for the non-modified proteome and phosphoproteome analysis, respectively. For these analyses, buffers A (99.9% H_2_O, 0.1% formic acid), B (99.9% ACN, 0.1% formic acid), C (100 mM ammonium acetate, 2% formic acid), and D (2 M ammonium acetate, 2% formic acid) were utilized. For each salt step, a 150-minute gradient program comprised of a 0–5 minute increase to the specified ammonium acetate concentration, 5–10 minutes hold, 10–14 minutes at 100% buffer A, 15–100 minutes 15–30% buffer B, 100-121 minutes 30–45% buffer B, 120–140 minutes 45–80% buffer B, 140–144 minutes 80% buffer B, and 145–150 minutes buffer A was employed. minutes buffer A was employed.

Eluted peptides were analyzed using a Thermo Scientific Q-Exactive Plus high-resolution quadrupole Orbitrap mass spectrometer, which was directly coupled to the HPLC. Data dependent acquisition was obtained using Xcalibur 4.0 software in positive ion mode with a spray voltage of 2.20 kV and a capillary temperature of 275 °C and an RF of 60. MS1 spectra were measured at a resolution of 70,000, an automatic gain control (AGC) of 3e6 with a maximum ion time of 100 ms and a mass range of 400-2000 m/z. Up to 15 MS2 were triggered at a resolution of 35,000 with a fixed first mass of 120 m/z for phosphoproteome and 120 m/z for proteome. An AGC of 1e5 with a maximum ion time of 50 ms, an isolation window of 1.3 m/z, and a normalized collision energy of 31. Charge exclusion was set to unassigned, 1, 5–8, and >8. MS1 that triggered MS2 scans were dynamically excluded for 45 or 25 s for phospho- and non-modified proteomes, respectively.

### Proteomics Data Analysis

The raw spectra were analyzed using MaxQuant version 1.6.14.0 (Tyanova *et al*, 2016). Spectra were searched using the Andromeda search engine in MaxQuant (Cox *et al*, 2011) against the Tair10 proteome file entitled “TAIR10_pep_20101214” that was downloaded from the TAIR website (https://www.arabidopsis.org/download_files/Proteins/TAIR10_protein_lists/TAIR10_pep_20101214) and was complemented with reverse decoy sequences and common contaminants by MaxQuant. Carbamidomethyl cysteine was set as a fixed modification while methionine oxidation and protein N-terminal acetylation were set as variable modifications. For the phosphoproteome “Phospho STY” was also set as a variable modification. The sample type was set to “Reporter Ion MS2” with “16plex TMT selected for both lysine and N-termini”. Digestion parameters were set to “specific” and “Trypsin/P;LysC”. Up to two missed cleavages were allowed. A false discovery rate, calculated in MaxQuant using a target-decoy strategy (Elias & Gygi, 2007), less than 0.01 at both the peptide spectral match and protein identification level was required. The ‘second peptide’ option identify co-fragmented peptides was not used. The match between runs feature of MaxQuant was not utilized.

Statistical analysis on the MaxQuant output was performed using the TMT-NEAT Analysis Pipeline (https://zenodo.org/badge/latestdoi/232925706) (Clark *et al*, 2021). Protein groups and phosphosites were categorized as differentially accumulating in each pairwise comparison if they had a *q-*value (i.e., adjusted p-value) < 0.1.

### Phosphoproteomic profiling

Phosphosites were classified by fold variation, having increased or decreased abundance and several lists were generated containing only the proteins with significant differences (q<0.1) compared to the Mock control for that specific time point and genotype. The locus codes for those genes were then compared to the Arabidopsis G-Signaling Interactome Database (AGIdb, http://bioin-folab.unl.edu/AGIdb) and additional interactions were identified by the Arabidopsis Interactions Viewer (http://bar.utoronto.ca/interactions2/). Network (**Fig 3E**) was created using the software Cytoscape.

### Gene ontology analysis

The lists generated by phosphoproteomic profiling were combined into two lists: proteins containing increased or decreased phosphosite abundance in Col-0. The two lists were submitted to PANTHER GO-SLIM molecular function analysis for an overrepresentation test and fold enrichment values were determined in comparison with the complete Arabidopsis Gene Database. Significant GO terms had a corrected *p-*value < 0.05.

### The Interactome construction and analysis

The Arabidopsis experimental protein-protein interaction (PPIE) network was curated from STRING (with experimental evidence) (Szklarczyk *et al*, 2015), Arabidopsis interactome map (AI-1MAIN) (Arabidopsis Interactome Mapping Consortium, 2011b), plant-pathogen immune network (PPIN-1 and 2) (Wessling *et al*, 2014; Mukhtar *et al*, 2011a), cell surface interactome (CSI) (Smakowska-Luzan *et al*, 2018), literature-curated interactions (LCI) (Lee *et al*, 2010), membrane-linked Interactome Database version 1 (MIND1) (Jones *et al*, 2014), EffectorK (González-Fuente *et al*, 2020) and BioGRID (Oughtred *et al*, 2019). The collective PPIE was used to extract the phosphorylated proteins interactors using in-house python scripts. The network was imported to Cytoscape for topology centrality analysis and visualization (Shannon *et al*, 2003). To identify the highly connected phosphorylated proteins in the PPIE network, we selected the degree centrality cutoff of ∼top 5% of nodes (degree ≥ 15; hub_15_) as described in (Mukhtar *et al*, 2011a; Ahmed *et al*, 2018a). The functional enrichment analysis was performed by Metascape (Zhou *et al*, 2019). The locus codes for those genes were then compared to the Arabidopsis G-Signaling Interactome Database (AGIdb, http://bioinfolab.unl.edu/AGIdb) and additional interactions were identified by the Arabidopsis Interactions Viewer (http://bar.utoronto.ca/interactions2/). Network (**Fig 2D**) was created using the software Cytoscape.

### Quantifying AtRGS1-YFP internalization

AtRGS1-YFP internalization was induced with 100 nM flg22 as described (Urano *et al*, 2012a; Fu *et al*, 2014b; Watkins *et al*, 2021). Briefly, 3-5 day-old etiolated Arabidopsis seeds expressing 35S:AtRGS1-YFP were treated with 1 nM flg22 2, 10, or 30 minutes respectively before imaging. Image acquisition was done on with a Zeiss LSM880 (Zeiss Microscopy, Oberkochen, Germany) with a C-Apochromat 40x/1.2NA water immersion objective. YFP excitation was at 514nm and emission collection 525-565nm. Emission collection was done with a GaAsP detector. Z-stack series was acquired at 0.5µm intervals between images with a pinhole yielding 1 Airy Unit. Image processing and RGS internalization measurements were done with Fiji ImageJ. Internalized YFP fluorescence was measured and subtracted from total YFP fluorescence of individual cells. Images were acquired on the hypocotyl epidermis 2-4 mm below the hypocotyls. Seedling exposure to light was minimized as much as was practical while imaging to avoid light induced internalization of AtRGS1.

*Pharmacological inhibition of baseline AtRGS1-YFP internalization and protein production in atbα-1*.

3-5 day-old etiolated seedlings were preincubated with 50 µM TyrA23 and MβCD 5 mM for 2 hours as previously described (Watkins *et al*, 2021). Seedlings were briefly washed with water and transferred to liquid growth media for 0, 2, 10, 30 or 60 minutes prior to imaging. For coincubation with endocytosis inhibitors and cycloheximide, seedlings were incubated with TyrA23 and MβCD for two hours prior to incubation with TyrA23, MβCD, and 200 µM cycloheximide for 1 hour. Imaging AtRGS1-YFP was done as described above.

### Quantification of AtRGS1-YFP abundance via immunoblot analyses

Seedlings of Col-0 and a*tbα-1* plants were grown in 1/2 MS for 7 days under low constant light. The seedlings were treated with the translation inhibitor cycloheximide at 200 uM for 30 minutes and, subsequently, with flg22 at 100 nM for 0, 3 and 15 minutes. Total protein was extracted as described by Liang et al., 2018. Protein levels were determined by Ponceau S staining and RGS1 levels were determined by probing with anti-GFP Tag polyclonal antibody (Invitrogen #A-11122). Bands were quantified using the software ImageJ and RuBisCO large chain (rbcL) was used as an endogenous control of total protein levels.

### Firefly Split Luciferase Assay

pCAMBIA/des/cLuc and pCAMBIA/des/nLuc (Lin et al., 2015) were used to generate the following plasmids: AtATBα-nLUC, and AtATBβ-nLUC. AtRGS1-nLUC, cLUC-AtGPA1, and AtAGB1-nLUC were previously reported (Watkins *et al*, 2021). pART27H-mCherry-AtAGG1 plasmid was obtained from Dr. Jose R Botella (University of Queensland, Brisbane, Australia). All plasmids were transformed into *A. tumefaciens* strain *GV3101*. nLUC and cLUC fusion partners were co-expressed in *N. benthamiana* leaves by agroinfiltration following protocols (Zhou *et al*, 2018). 48 hours after infiltration, 6mm leaf discs were collected to 96-well plate and 40µl 0.4mM D-luciferin was added to each well. Luminescence was measured by a spectraMax L microplate reader (Molecular Devices).

### Kinase inhibition assay

GST-WNK8 (0.5 ug), of GST-PP2A (5 ug) and/or His-RGS1 (10ug) were incubated in 15ul of kinase reaction buffer (5 mM Tris-HCl pH 7.5, 1 mM MgCl2, 0.4 mM ATP, 1mM PMSF) with a radio-labeled [gamma-32P]-ATP at 20°C for 4 hr. The samples were separated on a SDS-PAGE gel and exposed on an intensifying screen. The screen image and the Coomassie-stained gel are shown in **Fig 5**. The relative amounts of ^32^P were quantitated and provided as relative values in this figure.

### Luminol-based ROS analysis

flg22-induced ROS bursts were measured as described (Chung *et al*, 2014; Tunc-Ozdemir & Jones, 2017). Briefly, leaf discs from 5-week-old plants were placed singly into a 96-well plate with 250 μl of water per well. After overnight incubation, the water was replaced with 100 μl of reaction mix (17 μg/ml of Luminol (Sigma), 10 μg/ml of Horseradish Peroxidase (HRP; Sigma), and 100 nM flg22). Luminescence was measured immediately with 1 second integration and 2 minute intervals using a SpectraMax L (Molecular Device).

### Hypocotyl elongation assay

Assay performed as previously described (Jones *et al*, 2014). Briefly, seeds of Col-0 and null mutants were sterilized, then germinated on square plates with ½ x MS medium, pH 5.7, 0.8% (w/v) agar, supplemented with 1% (w/v) sucrose, and stratified for 4 days. Plates were light treated for four hours to induce germination, wrapped in aluminum foil, and covered to grow in the dark for 64 hours. Hypocotyls were imaged with a Nikon digital camera (D40) against a black background and quantified with Fiji ImageJ.

### Real Time RT-qPCR

RNA was extracted from seedlings grown on ¼ x MS medium, pH 5.7, 0.8% (w/v) agar, and 1% (w/v) sucrose for 1 week. cDNA library was prepared with a 1:1 mixture of oligo(dT) and random hexamer primers and Maxima Reverse Transcriptase enzyme (Thermo Scientific, Reference: EP0742). RT-qPCR analysis using this cDNA was performed on a MJ Research DNA Engine Opticon 2: Continuous Fluorescence Detector, using SYBR Green detection chemistry. Primers were ordered from Eton Bioscience Inc. and chosen to amplify a 200-300 bp amplicon 3’ of all t-DNA insertions (Appendix Table S2). TUBULIN4 primers [Fwd: AGAGGTTGACGAGCAGAT; Rev: ACCAATGAAAGTAGACGC] were used as an internal control to account for amount or RNA extracted across mutants. C(t) values were used to calculate fold induction results, which were normalized to Col-0.

### Statistical analyses

In Figs. 1, 3, 4, and 5A, one-way ANOVA with post-hoc Tukey HSD analysis was performed using GraphPad Prism 7. In Figure 4F, statistical differences between groups were established using unpaired t-test and, in Figure 5B, using a one-way ANOVA with post-hoc test using a Bonferroni approach. Gene ontology overrepresentation test was Fisher’s Exact with Bonferroni correction for multiple testing.

### Data availability

Raw proteomics data have been deposited on MassIVE with accession number MSV000085275 (ftp://massive.ucsd.edu/MSV000085275/).

FTP for reviewers: ftp://MSV000085275@massive.ucsd.edu

Password: “reviewer”

## Acknowledgements

This work was supported by grants from the NIGMS (GM065989) and NSF (MCB-1713880) to The Division of Chemical Sciences, Geosciences, and Biosciences, Office of Basic Energy Sciences of the US Department of Energy through the grant DE-FG02-05er15671 to A.M.J. funded the biochemical aspects of this project. J.M.W is supported by a NIGMS Institutional Research and Academic Career Development Award (GM000678). Research in J.W.W.’s laboratory is supported by NIH (GM120316), NSF (1759023 & 1818160), the ISU Plant Sciences, and USDA NIFA Hatch project IOW3808. N.M.C is supported by a USDA NIFA Postdoctoral Research Fellowship (2019-67012-29712). M.S.M is supported by NSF awards IOS-1557796 and IOS-2038872.

## Appendix (in order introduced in text)

1. Appendix Data Set 1: All phosphosites
2. Appendix Data Set 2: Protein abundance
3. Appendix Figure S1. Volcano plots of differential protein abundance and phosphosite expression.
4. Appendix Data Set S3: Modeling Data Supporting Main Figure 3
5. Appendix Figure S2. RGS1-YFP internalization in response to phosphatase inhibitors and phosphatase null mutations.
6. Appendix Data Set S4: Phosphorylated RLKs
7. Appendix Table S1: Published phosphoproteomes Conditions and Tissue overview
8. Appendix Table S2: Phosphatase mutant primer sequences
9. Appendix Data Set S5: TMT16 labeling Strategy

## APPENDIX

The following appendix was prepared per instructions to authors as follows:

1. A table of contents on the first page
2. Supplementary figures, text and simple tables and their legends (i.e. traditional Supplementary Information)
3. Use the nomenclature Appendix Figure S1, Appendix Table S1, Appendix Supplementary Methods etc. to ensure readers are not confused between Appendix figures and Expanded View figures
4. Reference these items in the manuscript text as: Appendix Figure S1, Appendix Table S1, Appendix Supplementary Methods
5. The Appendix PDF should be uploaded using the file type Expanded View File in our manuscript submission system.

**Appendix Table S1:** Published phosphoproteomes method overview.

|  | **Benschop, JJ.**<br>**2007** | **Nühse, TS.**<br>**2007** | **Rayapuram, N.**<br>**2014** | **Kohorn, BD.**<br>**2016** | **Watkins, JM.**<br>**2021**<br>**(this study)** |
| --- | --- | --- | --- | --- | --- |
| **Plant tissue** | Cultured cells | Cultured cells | Seedlings | Seedlings | 12-day-old roots |
| **Stimulus** | 1000 nM flg22 or<br>100 µg/ml xy-<br>lanase | 100 nM flg22 | 1000 nM flg22 | 50 µg/ml OG | 50 nM flg22 |
| **Time treated** | 10 min | 3-15 min | 15 min | 5 min | 3 and 15 min |

**Appendix Table S2:**
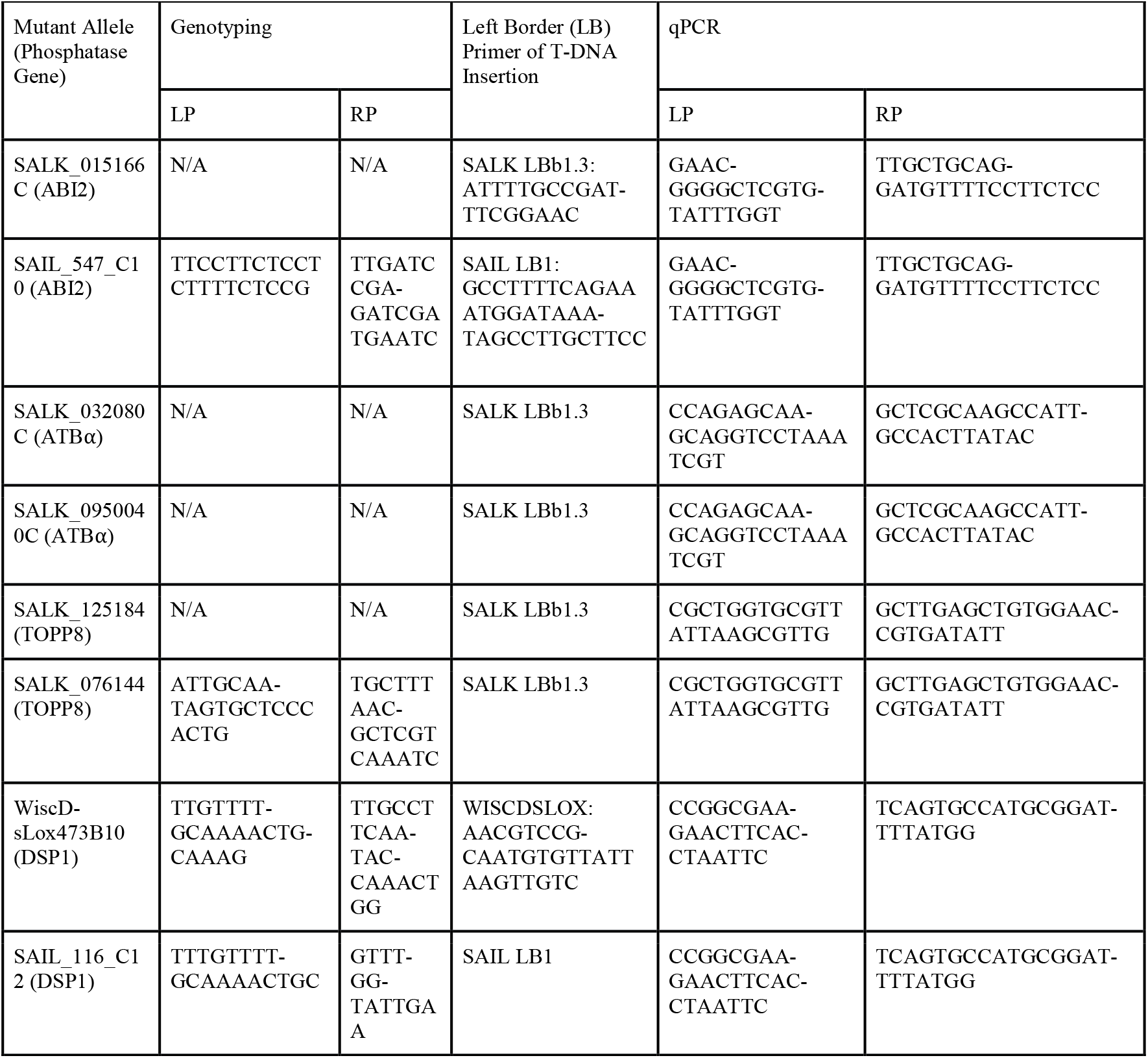
Phosphatase mutant primer sequences.

**Appendix Figure S1.**
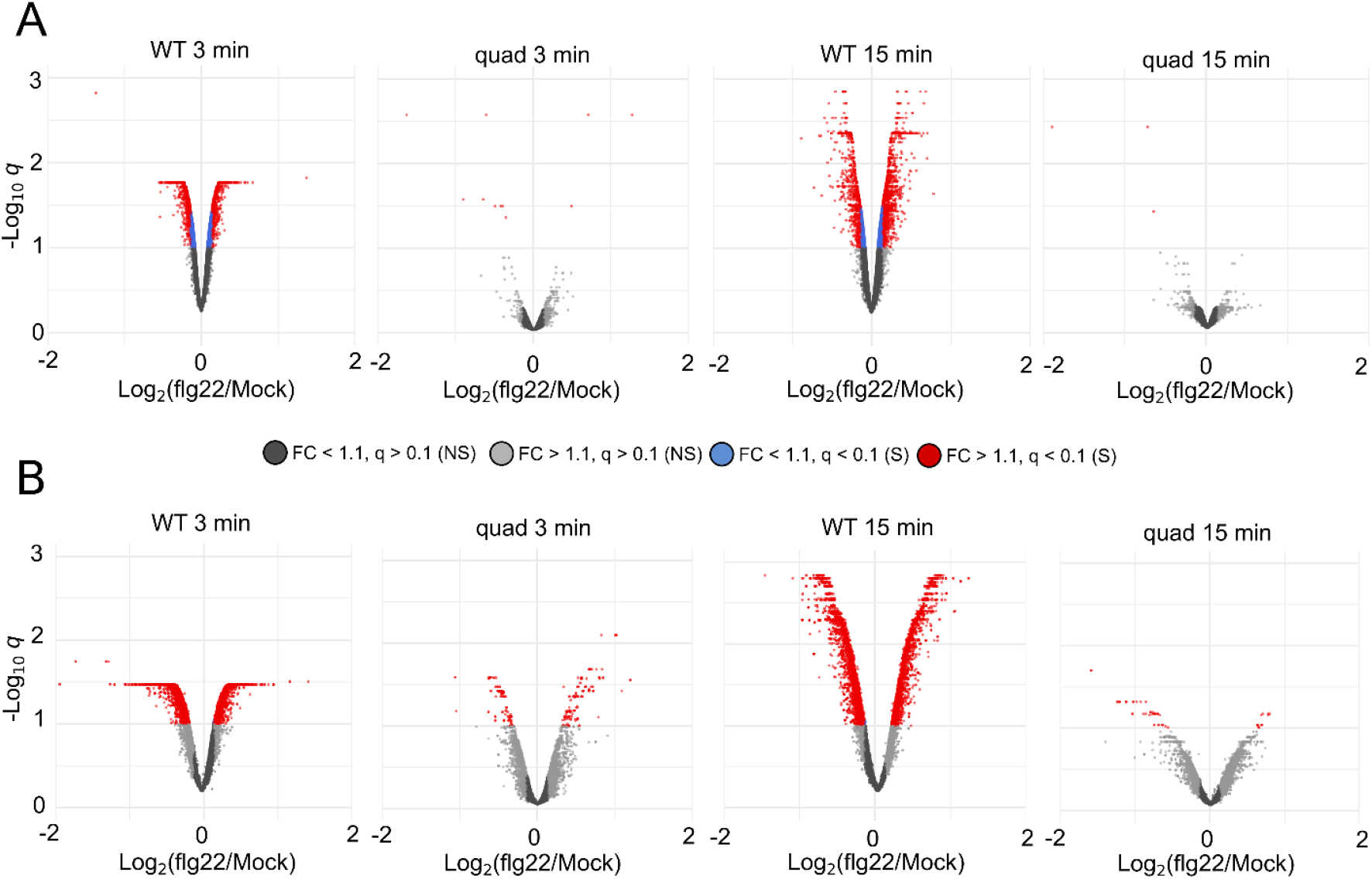
Volcano plots of differential protein abundance and phosphosite expression. A. Differential protein abundance B. Differential phosphosite response to flg22 treatment in WT and quad mutants after 3 or 15 min. Each dot represents one protein group (n = 8,918) or phosphosite (n = 24,468). x-axis is log2(flg22/mock). y-axis is -log10(q-value). Protein groups/phosphosites with a significant q-value (q < 0.1) are in blue (fold-change < 1.1) and red (fold change > 1.1). Protein groups/phosphosites with a non-significant q-value (q > 0.1) are in gray. S – significant, NS, not significant

**Appendix Figure S2.**
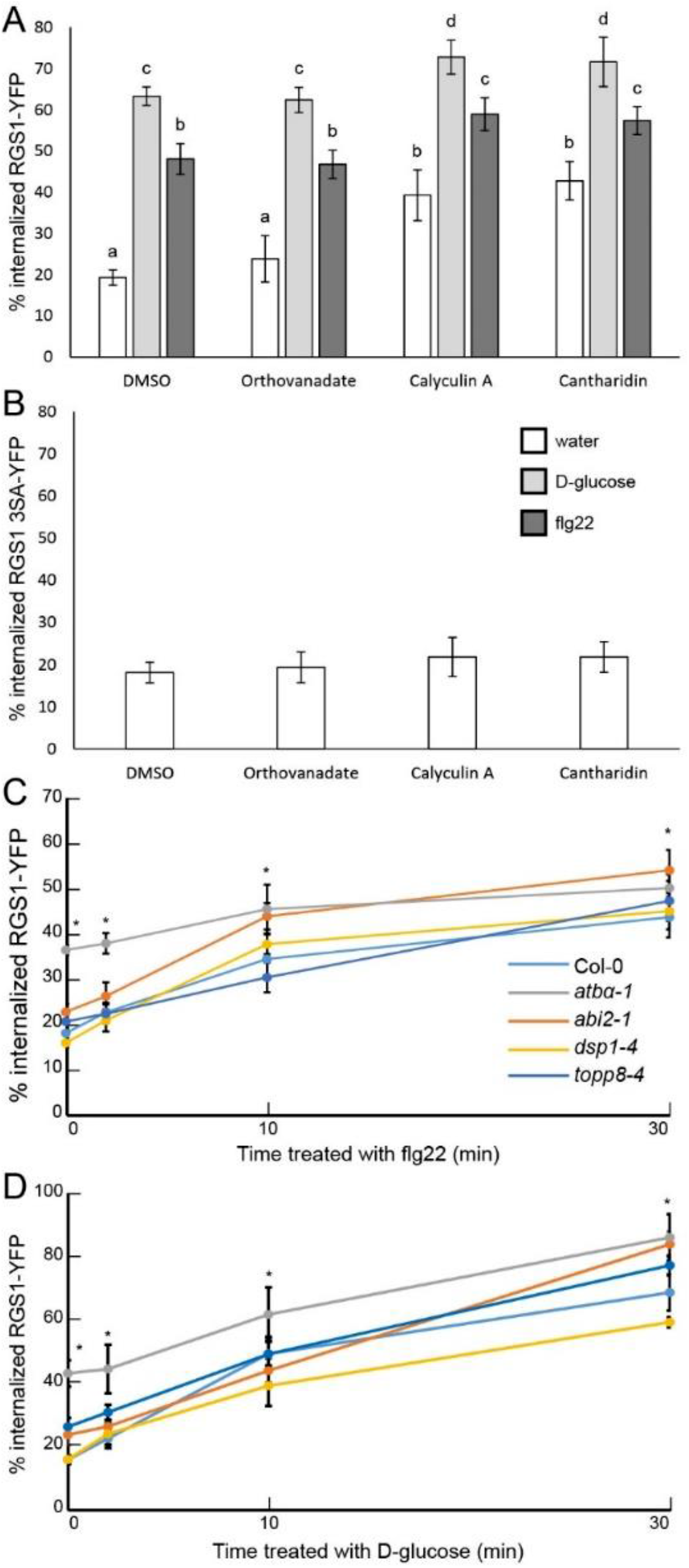
RGS1-YFP internalization in response to phosphatase inhibitors and phosphatase null mutations. A. D-glucose- or flg22-induced RGS1-YFP internalization after pretreatment with DMSO or phosphatase inhibitor: orthovanadate, calyculin A, and cantharidin for 2 hours. Means with different letters indicate significant difference (P < 0.05). Error bars represent CI. n = 15-30. B. RGS1-YFP internalization after treatment with DMSO or phosphatase inhibitor for 2 hours. Error bars represent CI. n = 28-35. C. flg22-induced RGS1-YFP internalization measured over time in Col-0 and null phosphatase mutants. * Represents statistical significance (P < 0.01) between Col-0 and atbα-1. Error bars represent CI. n=30 across three separate experimental replicates. D. D-glucose-induced RGS1-YFP internalization measured over time in Col-0 and null phosphatase mutants. * Represents statistical significance (P < 0.01) between Col-0 and atbα-1. Error bars represent CI. n=27-35 across three separate experimental replicates.

## References

Ahmed H, Howton TC, Sun Y, Weinberger N, Belkhadir Y & Mukhtar MS (2018a) Network biology discovers pathogen contact points in host protein-protein interactomes. Nat Commun 9: 2312

Ahmed H, Howton TC, Sun Y, Weinberger N, Belkhadir Y & Mukhtar MS (2018b) Network biology discovers pathogen contact points in host protein-protein interactomes. Nat Commun 9: 2312

Ahn I-P, Lee S-W & Suh S-C (2007) Rhizobacteria-Induced Priming in Arabidopsis Is Dependent on Ethylene, Jasmonic Acid, and NPR1. Mol Plant-Microbe Interactions® 20: 759–768

Altelaar AFM, Munoz J & Heck AJR (2013) Next-generation proteomics: towards an integrative view of proteome dynamics. Nat Rev Genet 14: 35–48

Arabidopsis Interactome Mapping Consortium (2011a) Evidence for network evolution in an Arabidopsis interactome map. Science 333: 601–607

Arabidopsis Interactome Mapping Consortium (2011b) Evidence for network evolution in an Arabidopsis interactome map. Science 333: 601–607

Bajsa J, Pan Z & Duke SO (2015) Cantharidin, a protein phosphatase inhibitor, strongly upregulates detoxification enzymes in the Arabidopsis proteome. J Plant Physiol 173: 33–40

Banbury DN, Oakley JD, Sessions RB & Banting G (2003) Tyrphostin A23 inhibits internalization of the transferrin receptor by perturbing the interaction between tyrosine motifs and the medium chain subunit of the AP-2 adaptor complex. J Biol Chem 278: 12022–12028

Beck M, Wyrsch I, Strutt J, Wimalasekera R, Webb A, Boller T & Robatzek S (2014) Expression patterns of FLAGELLIN SENSING 2 map to bacterial entry sites in plant shoots and roots. J Exp Bot 65: 6487– 6498

Benschop JJ, Mohammed S, O’Flaherty M, Heck AJR, Slijper M & Menke FLH (2007) Quantitative phosphoproteomics of early elicitor signaling in Arabidopsis. Mol Cell Proteomics MCP 6: 1198–1214

Chapman JM & Muday GK (2021) Flavonols modulate lateral root emergence by scavenging reactive oxygen species in Arabidopsis thaliana. J Biol Chem 296

Chen J-G, Willard FS, Huang J, Liang J, Chasse SA, Jones AM & Siderovski DP (2003) A seven-transmembrane RGS protein that modulates plant cell proliferation. Science 301: 1728–1731

Chinchilla D, Zipfel C, Robatzek S, Kemmerling B, Nürnberger T, Jones JDG, Felix G & Boller T (2007) A flagellin-induced complex of the receptor FLS2 and BAK1 initiates plant defence. Nature 448: 497– 500

Chung E-H, El-Kasmi F, He Y, Loehr A & Dangl JL (2014) A plant phosphoswitch platform repeatedly targeted by type III effector proteins regulates the output of both tiers of plant immune receptors. Cell Host Microbe 16: 484–494

Clark NM, Nolan TM, Wang P, Song G, Montes C, Valentine CT, Guo H, Sozzani R, Yin Y & Walley JW (2021) Integrated omics networks reveal the temporal signaling events of brassinosteroid response in Arabidopsis. Nat Commun 12: 5858

Cox J, Neuhauser N, Michalski A, Scheltema RA, Olsen JV & Mann M (2011) Andromeda: A Peptide Search Engine Integrated into the MaxQuant Environment. J Proteome Res 10: 1794–1805

Dai M, Zhang C, Kania U, Chen F, Xue Q, Mccray T, Li G, Qin G, Wakeley M, Terzaghi W, et al (2012) A PP6-Type Phosphatase Holoenzyme Directly Regulates PIN Phosphorylation and Auxin Efflux in Arabidopsis. Plant Cell 24: 2497–2514

Desaki Y, Miyata K, Suzuki M, Shibuya N & Kaku H (2018) Plant immunity and symbiosis signaling mediated by LysM receptors. Innate Immun 24: 92–100

Dhonukshe P, Aniento F, Hwang I, Robinson DG, Mravec J, Stierhof Y-D & Friml J (2007) Clathrin-mediated constitutive endocytosis of PIN auxin efflux carriers in Arabidopsis. Curr Biol CB 17: 520– 527

Elias JE & Gygi SP (2007) Target-decoy search strategy for increased confidence in large-scale protein identifications by mass spectrometry. Nat Methods 4: 207–214

Faris JD & Friesen TL (2020) Plant genes hijacked by necrotrophic fungal pathogens. Curr Opin Plant Biol 56: 74–80

Farkas I, Dombrádi V, Miskei M, Szabados L & Koncz C (2007) Arabidopsis PPP family of serine/threonine phosphatases. Trends Plant Sci 12: 169–176

Fu Y, Lim S, Urano D, Tunc-Ozdemir M, Phan NG, Elston TC & Jones AM (2014a) Reciprocal encoding of signal intensity and duration in a glucose-sensing circuit. Cell 156: 1084–1095

Fu Y, Lim S, Urano D, Tunc-Ozdemir M, Phan NG, Elston TC & Jones AM (2014b) Reciprocal encoding of signal intensity and duration in a glucose-sensing circuit. Cell 156: 1084–1095

Ghusinga KR, Elston TC & Jones AM (2021a) Towards resolution of a paradox in plant G-protein signaling. Plant Physiol (accepted)

Ghusinga KR, Paredes F, Jones AM & Colaneri A (2021b) Reported differences in the flg22 response of the null mutation of AtRGS1 correlates with fixed genetic variation in the background of Col-0 isolates. Plant Signal Behav 16: 1878685

González-Fuente M, Carrère S, Monachello D, Marsella BG, Cazalé A-C, Zischek C, Mitra RM, Rezé N, Cottret L, Mukhtar MS, et al (2020) EffectorK, a comprehensive resource to mine for Ralstonia, Xan-thomonas, and other published effector interactors in the Arabidopsis proteome. Mol Plant Pathol 21: 1257–1270

Gouveia BC, Calil IP, Machado JPB, Santos AA & Fontes EPB (2017) Immune Receptors and Co-receptors in Antiviral Innate Immunity in Plants. Front Microbiol 7: 2139

Halliwell B & Whiteman M (2004a) Measuring reactive species and oxidative damage in vivo and in cell culture: how should you do it and what do the results mean? Br J Pharmacol 142: 231–255

Halliwell B & Whiteman M (2004b) Measuring reactive species and oxidative damage in vivo and in cell culture: how should you do it and what do the results mean? Br J Pharmacol 142: 231–255

Heidari B, Nemie-Feyissa D, Kangasjärvi S & Lillo C (2013) Antagonistic Regulation of Flowering Time through Distinct Regulatory Subunits of Protein Phosphatase 2A. PLoS ONE 8: e67987

Hogrebe A, von Stechow L, Bekker-Jensen DB, Weinert BT, Kelstrup CD & Olsen JV (2018) Benchmarking common quantification strategies for large-scale phosphoproteomics. Nat Commun 9: 1045

Ilangumaran S & Hoessli DC (1998) Effects of cholesterol depletion by cyclodextrin on the sphingolipid microdomains of the plasma membrane. Biochem J 335: 433–440

Ishikawa A (2009) The Arabidopsis G-Protein β-Subunit Is Required for Defense Response against Agro-bacterium tumefaciens. Biosci Biotechnol Biochem 73: 47–52

Johnston CA, Taylor JP, Gao Y, Kimple AJ, Grigston JC, Chen J-G, Siderovski DP, Jones AM & Willard FS (2007) GTPase acceleration as the rate-limiting step in Arabidopsis G protein-coupled sugar signaling. Proc Natl Acad Sci 104: 17317–17322

Jones AM, Xuan Y, Xu M, Wang R-S, Ho C-H, Lalonde S, You CH, Sardi MI, Parsa SA, Smith-Valle E, et al (2014) Border Control—A Membrane-Linked Interactome of Arabidopsis. Science 344: 711–716

Jones JC, Jones AM, Temple BRS & Dohlman HG (2012) Differences in intradomain and interdomain motion confer distinct activation properties to structurally similar Gα proteins. Proc Natl Acad Sci U S A 109: 7275–7279

Jones JC, Temple BRS, Jones AM & Dohlman HG (2011) Functional Reconstitution of an Atypical G Protein Heterotrimer and Regulator of G Protein Signaling Protein (RGS1) from Arabidopsis thaliana. J Biol Chem 286: 13143–13150

Karampelias M, Neyt P, Groeve SD, Aesaert S, Coussens G, Rolčík J, Bruno L, Winne ND, Minnebruggen AV, Montagu MV, et al (2016) ROTUNDA3 function in plant development by phosphatase 2A-mediated regulation of auxin transporter recycling. Proc Natl Acad Sci 113: 2768–2773

Kataya ARA, Heidari B, Hagen L, Kommedal R, Slupphaug G & Lillo C (2015) Protein Phosphatase 2A Holoenzyme Is Targeted to Peroxisomes by Piggybacking and Positively Affects Peroxisomal β-Oxidation. Plant Physiol 167: 493–506

Klopffleisch K, Phan N, Augustin K, Bayne RS, Booker KS, Botella JR, Carpita NC, Carr T, Chen J-G, Cooke TR, et al (2011a) Arabidopsis G-protein interactome reveals connections to cell wall carbohydrates and morphogenesis. Mol Syst Biol 7: 532

Klopffleisch K, Phan N, Augustin K, Bayne RS, Booker KS, Botella JR, Carpita NC, Carr T, Chen J-G, Cooke TR, et al (2011b) Arabidopsis G-protein interactome reveals connections to cell wall carbohydrates and morphogenesis. Mol Syst Biol 7: 532

Kohorn BD, Hoon D, Minkoff BB, Sussman MR & Kohorn SL (2016) Rapid Oligo-Galacturonide Induced Changes in Protein Phosphorylation in Arabidopsis. Mol Cell Proteomics MCP 15: 1351–1359

Lee K, Thorneycroft D, Achuthan P, Hermjakob H & Ideker T (2010) Mapping plant interactomes using literature curated and predicted protein-protein interaction data sets. Plant Cell 22: 997–1005

Li B, Tunc-Ozdemir M, Urano D, Jia H, Werth EG, Mowrey DD, Hicks LM, Dokholyan NV, Torres MP & Jones AM (2018) Tyrosine phosphorylation switching of a G protein. J Biol Chem 293: 4752–4766

Li R, Liu P, Wan Y, Chen T, Wang Q, Mettbach U, Baluška F, Šamaj J, Fang X, Lucas WJ, et al (2012) A Membrane Microdomain-Associated Protein, Arabidopsis Flot1, Is Involved in a Clathrin-Independent Endocytic Pathway and Is Required for Seedling Development. Plant Cell 24: 2105–2122

Liang X, Ding P, Lian K, Wang J, Ma M, Li L, Li L, Li M, Zhang X, Chen S, et al (2016) Arabidopsis heterotrimeric G proteins regulate immunity by directly coupling to the FLS2 receptor. eLife 5: e13568

Liang X, Ma M, Zhou Z, Wang J, Yang X, Rao S, Bi G, Li L, Zhang X, Chai J, et al (2018) Ligand-triggered derepression of Arabidopsis heterotrimeric G proteins coupled to immune receptor kinases. Cell Res 28: 529–543

Liao K-L, Melvin CE, Sozzani R, Jones RD, Elston TC & Jones AM (2017) Dose-Duration Reciprocity for G protein activation: Modulation of kinase to substrate ratio alters cell signaling. PLOS ONE 12: e0190000

Lorek J, Griebel T, Jones AM, Kuhn H & Panstruga R (2013) The role of Arabidopsis heterotrimeric G-protein subunits in MLO2 function and MAMP-triggered immunity. Mol Plant-Microbe Interact MPMI 26: 991–1003

Lu D, Wu S, Gao X, Zhang Y, Shan L & He P (2010) A receptor-like cytoplasmic kinase, BIK1, associates with a flagellin receptor complex to initiate plant innate immunity. Proc Natl Acad Sci 107: 496– 501

McAlister GC, Huttlin EL, Haas W, Ting L, Jedrychowski MP, Rogers JC, Kuhn K, Pike I, Grothe RA, Blethrow JD, et al (2012) Increasing the Multiplexing Capacity of TMTs Using Reporter Ion Isotopologues with Isobaric Masses. Anal Chem 84: 7469–7478

Merlot S, Gosti F, Guerrier D, Vavasseur A & Giraudat J (2001) The ABI1 and ABI2 protein phosphatases 2C act in a negative feedback regulatory loop of the abscisic acid signalling pathway. Plant J Cell Mol Biol 25: 295–303

Merlot S, Leonhardt N, Fenzi F, Valon C, Costa M, Piette L, Vavasseur A, Genty B, Boivin K, Müller A, et al (2007) Constitutive activation of a plasma membrane H+-ATPase prevents abscisic acid-mediated stomatal closure. EMBO J 26: 3216–3226

Millet YA, Danna CH, Clay NK, Songnuan W, Simon MD, Werck-Reichhart D & Ausubel FM (2010) Innate Immune Responses Activated in Arabidopsis Roots by Microbe-Associated Molecular Patterns. Plant Cell 22: 973–990

Mishra B, Kumar N & Mukhtar MS (2019) Systems Biology and Machine Learning in Plant–Pathogen Interactions. Mol Plant-Microbe Interactions® 32: 45–55

Mishra B, Kumar N & Mukhtar MS (2021a) Network biology to uncover functional and structural properties of the plant immune system. Curr Opin Plant Biol 62: 102057

Mishra B, Kumar N & Mukhtar MS (2021b) Network biology to uncover functional and structural properties of the plant immune system. Curr Opin Plant Biol 62: 102057

Mishra B, Sun Y, Ahmed H, Liu X & Mukhtar MS (2017a) Global temporal dynamic landscape of pathogen-mediated subversion of Arabidopsis innate immunity. Sci Rep 7: 7849

Mishra B, Sun Y, Ahmed H, Liu X & Mukhtar MS (2017b) Global temporal dynamic landscape of pathogen-mediated subversion of Arabidopsis innate immunity. Sci Rep 7: 7849

Mott GA, Smakowska-Luzan E, Pasha A, Parys K, Howton TC, Neuhold J, Lehner A, Grünwald K, StoltBergner P, Provart NJ, et al (2019) Map of physical interactions between extracellular domains of Arabidopsis leucine-rich repeat receptor kinases. Sci Data 6: 190025

Mukhtar MS, Carvunis A-R, Dreze M, Epple P, Steinbrenner J, Moore J, Tasan M, Galli M, Hao T, Nishimura MT, et al (2011a) Independently evolved virulence effectors converge onto hubs in a plant immune system network. Science 333: 596–601

Mukhtar MS, Carvunis A-R, Dreze M, Epple P, Steinbrenner J, Moore J, Tasan M, Galli M, Hao T, Nishimura MT, et al (2011b) Independently Evolved Virulence Effectors Converge onto Hubs in a Plant Immune System Network. Science 333: 596–601

Nühse TS, Bottrill AR, Jones AME & Peck SC (2007) Quantitative phosphoproteomic analysis of plasma membrane proteins reveals regulatory mechanisms of plant innate immune responses. Plant J Cell Mol Biol 51: 931–940

Ohtani Y, Irie T, Uekama K, Fukunaga K & Pitha J (1989) Differential effects of alpha-, beta- and gamma-cyclodextrins on human erythrocytes. Eur J Biochem 186: 17–22

Oughtred R, Stark C, Breitkreutz B-J, Rust J, Boucher L, Chang C, Kolas N, O’Donnell L, Leung G, McAdam R, et al (2019) The BioGRID interaction database: 2019 update. Nucleic Acids Res 47: D529–D541

Patharkar OR, Gassmann W & Walker JC (2017) Leaf shedding as an anti-bacterial defense in Arabidopsis cauline leaves. PLOS Genet 13: e1007132

Rayapuram N, Bigeard J, Alhoraibi H, Bonhomme L, Hesse A-M, Vinh J, Hirt H & Pflieger D (2018) Quantitative Phosphoproteomic Analysis Reveals Shared and Specific Targets of Arabidopsis Mitogen-Activated Protein Kinases (MAPKs) MPK3, MPK4, and MPK6. Mol Cell Proteomics MCP 17: 61–80

Rayapuram N, Bonhomme L, Bigeard J, Haddadou K, Przybylski C, Hirt H & Pflieger D (2014) Identification of novel PAMP-triggered phosphorylation and dephosphorylation events in Arabidopsis thaliana by quantitative phosphoproteomic analysis. J Proteome Res 13: 2137–2151

Romá-Mateo C, Sacristán-Reviriego A, Beresford NJ, Caparrós-Martín JA, Culiáñez-Macià FA, Martín H, Molina M, Tabernero L & Pulido R (2011) Phylogenetic and genetic linkage between novel atypical dual-specificity phosphatases from non-metazoan organisms. Mol Genet Genomics MGG 285: 341–354

Segonzac C, Macho AP, Sanmartín M, Ntoukakis V, Sánchez-Serrano JJ & Zipfel C (2014) Negative control of BAK1 by protein phosphatase 2A during plant innate immunity. EMBO J 33: 2069–2079

Shannon P, Markiel A, Ozier O, Baliga NS, Wang JT, Ramage D, Amin N, Schwikowski B & Ideker T (2003) Cytoscape: a software environment for integrated models of biomolecular interaction networks. Genome Res 13: 2498–2504

Smakowska-Luzan E, Mott GA, Parys K, Stegmann M, Howton TC, Layeghifard M, Neuhold J, Lehner A, Kong J, Grünwald K, et al (2018) An extracellular network of Arabidopsis leucine-rich repeat receptor kinases. Nature 553: 342–346

Song G, Brachova L, Nikolau BJ, Jones AM & Walley JW (2018a) Heterotrimeric G-Protein-Dependent Proteome and Phosphoproteome in Unstimulated Arabidopsis Roots. Proteomics 18: e1800323

Song G, Hsu PY & Walley JW (2018b) Assessment and Refinement of Sample Preparation Methods for Deep and Quantitative Plant Proteome Profiling. Proteomics 18: e1800220

Stubbs MD, Tran HT, Atwell AJ, Smith CS, Olson D & Moorhead GBG (2001) Purification and properties of Arabidopsis thaliana type 1 protein phosphatase (PP1). Biochim Biophys Acta BBA - Protein Struct Mol Enzymol 1550: 52–63

Sun Y, Li L, Macho AP, Han Z, Hu Z, Zipfel C, Zhou J-M & Chai J (2013) Structural basis for flg22-induced activation of the Arabidopsis FLS2-BAK1 immune complex. Science 342: 624–628

Szklarczyk D, Franceschini A, Wyder S, Forslund K, Heller D, Huerta-Cepas J, Simonovic M, Roth A, Santos A, Tsafou KP, et al (2015) STRING v10: protein-protein interaction networks, integrated over the tree of life. Nucleic Acids Res 43: D447–452

Teixeira RM, Ferreira MA, Raimundo GAS, Loriato VAP, Reis PAB & Fontes EPB (2019) Virus perception at the cell surface: revisiting the roles of receptor-like kinases as viral pattern recognition receptors. Mol Plant Pathol 20: 1196–1202

Thung L, Trusov Y, Chakravorty D & Botella JR (2012) Gγ1+Gγ2+Gγ3=Gβ: the search for heterotrimeric G-protein γ subunits in Arabidopsis is over. J Plant Physiol 169: 542–545

Toorn M aan den, Huijbers MME, Vries SC de & Mierlo CPM van (2012) The Arabidopsis thaliana SERK1 Kinase Domain Spontaneously Refolds to an Active State In Vitro. PLOS ONE 7: e50907

Tsvetanova NG, Trester-Zedlitz M, Newton BW, Peng GE, Johnson JR, Jimenez-Morales D, Kurland AP, Krogan NJ & von Zastrow M (2021) Endosomal cAMP production broadly impacts the cellular phosphoproteome. J Biol Chem 297: 100907

Tunc-Ozdemir M & Jones AM (2017) Ligand-induced dynamics of heterotrimeric G protein-coupled receptor-like kinase complexes. PLOS ONE 12: e0171854

Tunc-Ozdemir M, Li B, Jaiswal DK, Urano D, Jones AM & Torres MP (2017) Predicted Functional Implications of Phosphorylation of Regulator of G Protein Signaling Protein in Plants. Front Plant Sci 8

Tunc-Ozdemir M, Urano D, Jaiswal DK, Clouse SD & Jones AM (2016) Direct Modulation of Heterotrimeric G Protein-coupled Signaling by a Receptor Kinase Complex. J Biol Chem 291: 13918–13925

Tyanova S, Temu T & Cox J (2016) The MaxQuant computational platform for mass spectrometry-based shotgun proteomics. Nat Protoc 11: 2301–2319

Tyburski J, Dunajska K & Tretyn A (2009) Reactive oxygen species localization in roots of Arabidopsis thaliana seedlings grown under phosphate deficiency. Plant Growth Regul 59: 27–36

Urano D, Maruta N, Trusov Y, Stoian R, Wu Q, Liang Y, Jaiswal DK, Thung L, Jackson D, Botella JR, et al (2016) Saltational evolution of the heterotrimeric G protein signaling mechanisms in the plant kingdom. Sci Signal 9: ra93

Urano D, Phan N, Jones JC, Yang J, Huang J, Grigston J, Philip Taylor J & Jones AM (2012a) Endocytosis of the seven-transmembrane RGS1 protein activates G-protein-coupled signalling in Arabidopsis. Nat Cell Biol 14: 1079–1088

Urano D, Phan N, Jones JC, Yang J, Huang J, Grigston J, Taylor JP & Jones AM (2012b) Endocytosis of the seven-transmembrane RGS1 protein activates G-protein-coupled signalling in Arabidopsis. Nat Cell Biol 14: 1079–1088

Watkins JM, Chapman JM & Muday GK (2017) Abscisic Acid-Induced Reactive Oxygen Species Are Modulated by Flavonols to Control Stomata Aperture. Plant Physiol 175: 1807–1825

Watkins JM, Hechler PJ & Muday GK (2014) Ethylene-Induced Flavonol Accumulation in Guard Cells Suppresses Reactive Oxygen Species and Moderates Stomatal Aperture. Plant Physiol 164: 1707– 1717

Watkins JM, Ross-Elliott TJ, Shan X, Lou F, Dreyer B, Tunc-Ozdemir M, Jia H, Yang J, Oliveira CC, Wu L, et al (2021) Differential regulation of G protein signaling in Arabidopsis through two distinct pathways that internalize AtRGS1. Sci Signal 14: eabe4090

Wessling R, Epple P, Altmann S, He Y, Yang L, Henz SR, McDonald N, Wiley K, Bader KC, Gläßer C, et al (2014) Convergent targeting of a common host protein-network by pathogen effectors from three kingdoms of life. Cell Host Microbe 16: 364–375

Wolfenstetter S, Chakravorty D, Kula R, Urano D, Trusov Y, Sheahan MB, McCurdy DW, Assmann SM, Jones AM & Botella JR (2015) Evidence for an unusual transmembrane configuration of AGG3, a Class C Gγ Subunit, of Arabidopsis. Plant J Cell Mol Biol 81: 388–398

Xu Y, Xing Y, Chen Y, Chao Y, Lin Z, Fan E, Yu JW, Strack S, Jeffrey PD & Shi Y (2006) Structure of the protein phosphatase 2A holoenzyme. Cell 127: 1239–1251

Xue J, Gong B-Q, Yao X, Huang X & Li J-F (2020) BAK1-mediated phosphorylation of canonical G protein alpha during flagellin signaling in Arabidopsis. J Integr Plant Biol 62: 690–701

Yang Z, Xing J, Wang L, Liu Y, Qu J, Tan Y, Fu X, Lin Q, Deng H & Yu F (2020) Mutations of two FERONIA-like receptor genes enhance rice blast resistance without growth penalty. J Exp Bot 71: 2112– 2126

Yemets A, Sheremet Y, Vissenberg K, Van Orden J, Verbelen J-P & Blume YB (2008) Effects of tyrosine kinase and phosphatase inhibitors on microtubules in Arabidopsis root cells. Cell Biol Int 32: 630–637

Yu Y, Chakravorty D & Assmann SM (2018) The G Protein β-Subunit, AGB1, Interacts with FERONIA in RALF1-Regulated Stomatal Movement. Plant Physiol 176: 2426–2440

Zhou Y, Zhou B, Pache L, Chang M, Khodabakhshi AH, Tanaseichuk O, Benner C & Chanda SK (2019) Metascape provides a biologist-oriented resource for the analysis of systems-level datasets. Nat Commun 10: 1523

Zhou Z, Bi G & Zhou J-M (2018) Luciferase Complementation Assay for Protein-Protein Interactions in Plants. Curr Protoc Plant Biol 3: 42–50

